# BMAL1 and MEX3A co-regulate intestinal stem cell succession

**DOI:** 10.1101/2022.03.08.483537

**Authors:** Fang-Pei Chang, Cheng-Kai Wang, Yu-Chi Chou, Tzu-Jou Chen, Ya-Wen Wu, Samyuktha Sridharan, Pang-Hung Hsu, Wendy W. Hwang-Verslues

## Abstract

Chemotherapeutic agents, such as 5-fluorouracil (5-FU), often induce intestinal mucositis due to toxicity to rapidly dividing intestinal stem cells (ISCs). Drug delivery at different times of day can alter 5-FU toxicity; however, the underlying biology is unclear. We found that homeostasis and succession between two ISC types, fast proliferating Lgr5^+^ crypt base columnar (CBC) cells and stress-resistant +4 cells (Bmi1^+^), are controlled by the circadian clock transcription factor BMAL1 and the BMAL1-regulated RNA-binding protein MEX3A via both direct transcriptional regulation and post-transcriptional control of *Lgr5* mRNA stability. *Bmal1* knockout in Lgr5^+^ CBCs reduced MEX3A expression and CBC numbers but increased Bmi1 expression in the crypts. Timing 5-FU delivery when crypt cells had low *Lgr5* but peak level of *Bmi1* protected ISCs from apoptosis. Together, these findings identify a novel role of BMAL1 in ISC homeostasis and provide a biological explanation for chronotherapeutic chemoprotection.

## INTRODUCTION

Chemotherapy is currently the most effective cancer treatment. However, these anti-cancer drugs target both fast proliferating cancer cells and rapidly dividing normal cells, such as intestinal epithelial cells, and therefore result in adverse effects. For example, 5-fluorouracil (5-FU) is the first-line treatment for colorectal cancer. However, patients often suffer from intestinal mucositis due to damage to intestinal epithelial cells. Mitigation of such side effects while maintaining anti-cancer effectiveness of chemotherapy has been a major challenge in cancer management. Many factors contributing to drug transport and metabolism are regulated by circadian oscillators (Dae Wook Kim, 2020). Moreover, many proteins identified as druggable targets by the US Food and Drug Administration are encoded by genes with rhythmic expression (Rijo-Ferreira and Takahashi, 2019). Thus, developing chrono-modulated chemotherapy by administrating drugs at specific times of day may provide an opportunity to maximize the efficacy of the treatment and minimize side effects (Levi et al., 2010). To precisely deliver chemotherapeutic agents at the right time, it is essential to understand the biological rhythms and the stress response specifically in the key affected cells.

A healthy intestine is essential for food digestion, nutrient extraction, absorption and waste removal. The intestinal epithelium is organized into crypt-villus structures. In the crypts, there are two types of intestinal stem cells (ISCs) wedged between Paneth cells which protect and provide growth factors to the ISCs (Gehart and Clevers, 2019). One type of ISCs is LGR5-expressing crypt base columnar stem (CBC) cells which sit at position +1 to +3 at the crypt base between Paneth cells. The other ISC type is ‘+4’ cells which express BMI1, HOPX and LRIG1 (Gehart and Clevers, 2019). Proliferation of LGR5^+^ CBCs drives the rapid turnover and regeneration of intestinal epithelium (Barker et al., 2007; van der Flier and Clevers, 2009). However, the fast-proliferating LGR5^+^ CBCs are sensitive to cytotoxic injury resulting from chemotherapy or irradiation (IR). Depletion of LGR5^+^ CBCs by IR resulted in severe crypt loss and irregular villus structure (Metcalfe et al., 2014), indicating that replenishment of LGR5^+^ CBCs is essential for the intestinal epithelium to regenerate properly after injury. In contrast, BMI1^+^ +4 cells are more resistant to IR (Yan et al., 2012). It has been suggested that BMI1^+^ +4 cells can reprogram into LGR5-expressing cells to replenish the crypt base when LGR5^+^ CBCs are depleted due to intestinal injury (Tian et al., 2011). However, it is not known what regulates the homeostasis and succession between these two types of ISCs. A few recent studies have suggested that circadian rhythm affects ISC proliferation and tissue regeneration (Matsu-Ura et al., 2016; Stokes et al., 2017). However, whether the core circadian machinery regulates LGR5 and BMI1 expression and contributes to the succession between LGR5^+^- and BMI1^+^-ISCs remain to be determined.

Among the core clock genes, BMAL1 is the key transcription factor (Bunger et al., 2000). BMAL1 forms heterodimers with CLOCK (Hogenesch et al., 1998) to transcriptionally regulate other genes required for the complex network of transcription-translation feedback loops (TTFL) that drive circadian rhythms. In the gastrointestinal (GI) tract, it has been shown that BMAL1 deletion inhibited drug export (Yu et al., 2019), prevented ghrelin secretion (Laermans et al., 2015), and promoted glucose uptake (Sussman et al., 2019). These observations demonstrated an indispensable role of BMAL1 in maintaining GI tract function. BMAL1 appeared to regulate the timely division of intestinal cells as depletion of BMAL1 abolished oscillation of WNT3A, a critical niche component for maintaining LGR5^+^ CBC proliferation, and resulted in disruption of circadian rhythm and cell cycle coupling (Matsu-Ura et al., 2016). In addition, BMAL1 was found to contribute to the inflammation response and cell proliferation during intestinal regeneration (Stokes et al., 2017). Despite mounting evidence suggesting that BMAL1 may be essential to intestinal cell proliferation and regeneration, how BMAL1 regulates the homeostasis and succession of ISCs remains unclear.

In any given mammalian cell or tissue, approximately 5-20% of the transcripts show circadian oscillations (Takahashi, 2017), and approximately 35% of the oscillating transcripts are modulated by post-transcriptional control (Wang et al., 2018). RNA-binding proteins (RBPs) regulate various aspects of RNA fate and function (Garneau et al., 2007; Hentze et al., 2018). Several RBPs participate in circadian regulation (Kim et al., 2007; Lee et al., 2014; Lee et al., 2012). In the intestine, RBPs, including MEX3A, MSI1, HUR and IMP1, were found to facilitate intestinal development and repair, as well as maintenance of normal mucosa structures (Chatterji and Rustgi, 2018; Pereira et al., 2020). It is possible that the circadian machinery incorporates RBP-mediated post-transcriptional regulation to maintain intestinal homeostasis.

MEX3A expression has been used to identify a subset of slowly dividing LGR5^+^ CBCs (Barriga et al., 2017). Under chemotherapy-induced intestinal damage, these MEX3A^high^ cells contributed to intestinal epithelium repair and homeostasis (Barriga et al., 2017). In addition, MEX3A knockout decreased the number of LGR5^+^ CBCs in the duodenum and delayed organoid formation (Pereira et al., 2020), indicating that MEX3A is required for survival and stemness functions of LGR5^+^ CBCs. However, the underlying mechanism of MEX3A effect on LGR5^+^ CBCs is unknown.

We found that BMAL1 and MEX3A act together to regulate *Lgr5* expression and maintain the homeostasis between LGR5^+^ CBCs and BMI1^+^ +4 cells in intestinal crypts. In this mechanism, BMAL1 upregulated expression of both *Lgr5* and *Mex3a* and while Mex3a further enhanced Lgr5 expression by directly binding and stabilizing *Lgr5* mRNA. We then used this mechanistic insight to select a circadian time where 5-FU application caused minimal level of ISC apoptosis. These findings can be used to design chrono-chemotherapy strategies to increase drug efficacy and decrease side effects.

## RESULTS

### Rhythmic expression of *Mex3a*, *Lgr5* and *Bmi1* are correlated with BMAL1 ultradian expression in the crypts

We observed a 24-hour oscillation of the core clock gene *Bmal1* in the duodenum of WT *C57BL/6JNarl* mice kept under LD condition (Figure 1A). *Bmal1* mRNA decreased to its lowest level before the subjective dusk (zeitgeber time (ZT), ZT0 corresponds to lights on; ZT12 corresponds to lights off) and reached the peak level before the subjective dawn (Figure 1A), consistent with its expression pattern in the suprachiasmatic nucleus (SCN) (Honma et al., 1998) and other peripheral tissues (Welz et al., 2019; Yamamoto et al., 2004). BMAL1 protein expression was high during the light phase (peak at ZT5), but low during the dark period in the duodenum epithelium (Figure 1B). Surprisingly, immunohistochemistry (IHC) of duodenum crypts showed an ultradian oscillation of BMAL1 protein which peaked at ZT5 and ZT17 (Figure 1C; (note that an ultradian oscillation is defined as any oscillation with period shorter than 24 h; (Aschoff, 1981)). Moreover, RNAscope analyses revealed that gene expression of CBC markers, *Mex3a* and *Lgr5*, also oscillated (Figure 1C). *Mex3a* mRNA oscillation coincided with BMAL1 protein expression (Figure 1C and 1D). In contrast, *Lgr5* expression showed an antiphase pattern compared to the BMAL1 protein level (Figure 1C and 1E). Interestingly, the mRNA of the +4 cell marker, *Bmi1*, showed a 24 h oscillation which gradually increased from ZT9 to ZT17 when BMAL1 expression was low, and decreased when BMAL1 level was high in the crypt cells (Figure 1C and 1F). The unexpected ultradian oscillation of BMAL1 in the crypt cells and its correlation with the expression patterns of both CBC and +4 ISC markers suggest that ISC homeostasis may depend on BMAL1-mediated rhythmic regulation.

**Figure 1.**
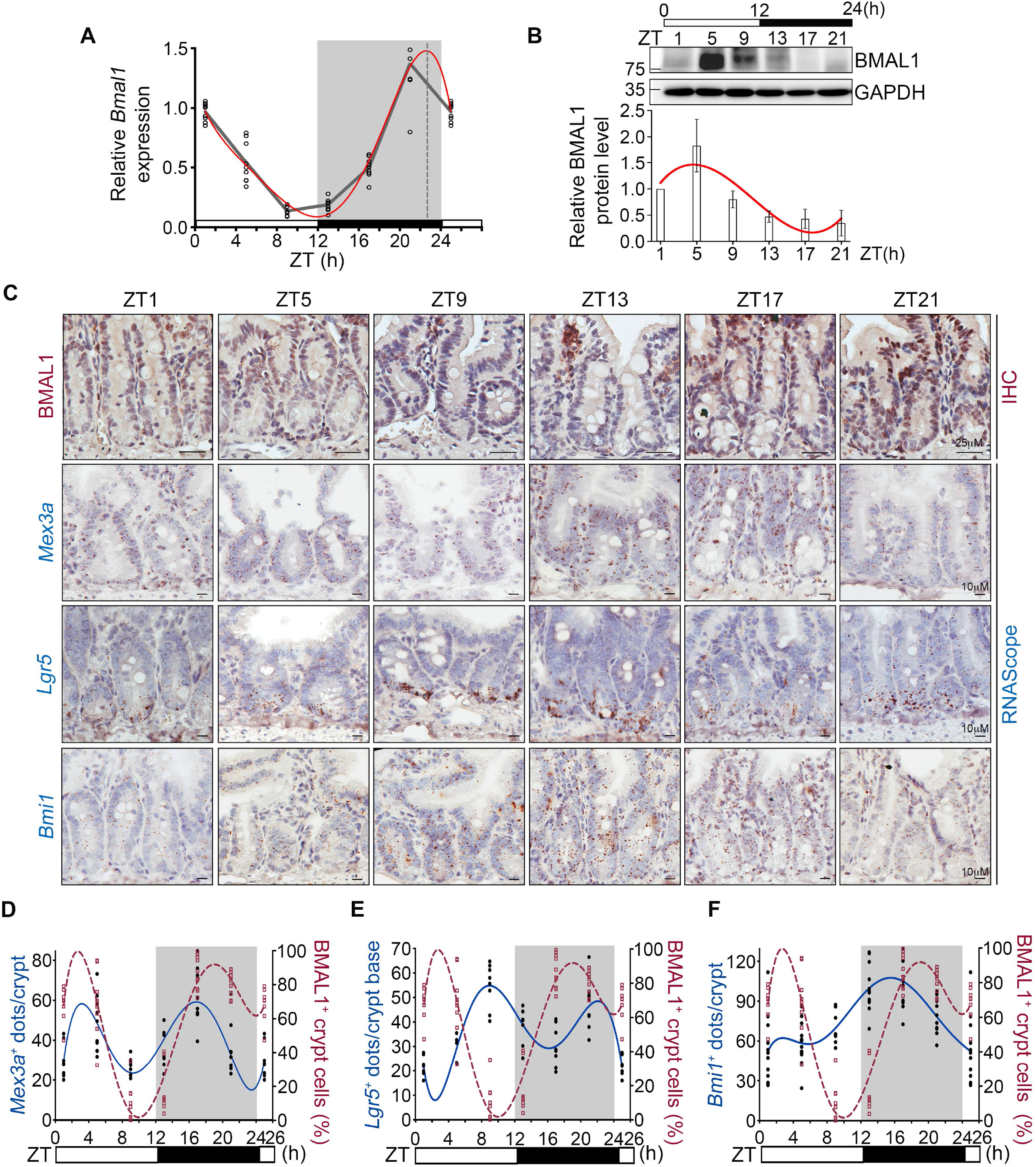
Oscillation of *Mex3a*, *Lgr5* and *Bmi1* in crypt cells is associated with BMAL1 ultradian expression. (A) qRT-PCR analysis of *Bmal1* level in WT *C57BL/6JNarl* (B6) duodenum harvested every 4 hours under LD condition. Curve-fitting results of relative *Bmal1* mRNA level are analyzed by nonlinear regression (red line). Three independent experiments were performed and data are means ± SD from one representative experiment (n =3). (B) Immunoblot of BMAL1 expression in WT B6 duodenum harvested every 4 hours under LD condition. GAPDH was used as a loading control. Blots shown are from one representative experiment of three independent experiment. (C) Representative images of IHC staining to detect BMAL1 protein and RNAscope of *Mex3a*, *Lgr5*, and *Bmi1* mRNA levels in WT B6 duodenum at the indicated ZT time points. Scale bars, 10 μM. (n ≧ 2 mice per time point) (D-F) Quantification of IHC and RNAscope analysis of BMAL1, *Mex3a*, *Lgr5*, and *Bmi1* expression in B6 crypts. The prevalence of BMAL1^+^ cells in crypts from IHC analysis (red dashed line) was quantified using the QuPath software. Cells with a mean DAB OD value greater than 0.2 are classified as BMAL1^+^ cells. Quantification of *Mex3a* (D), *Lgr5* (E) and *Bmi1* (F) mRNA positive dots (blue line) was performed using the trainable weka segmentation classifier in Fiji software. (n = ≧ 2 mice per time point, ≥ 50 crypts/n)

### BMAL1 is essential for maintaining LGR5 expression and ISC homeostasis in the crypts

It is generally accepted that loss of *Bmal1* expression in a specific tissue causes loss of circadian rhythm in that tissue (Dyar et al., 2014; Lamia et al., 2008). To determine the role of BMAL1 in LGR5^+^ CBCs, we established a tamoxifen (TAM) induced-LGR5-expressing cell specific *Bmal1* knockout mouse model [*Lgr5-Cre^+^;Bmal1^fl/fl^* (*LC^+^B^fl/fl^*)] by crossing *B6.129P2-Lgr5^tm1(cre/ERT2)Cle^/J* and *B6.129S4(Cg)-Arntl^tm1Weit^/J* mice (Figure 2A). After TAM-induced *Bmal1* knockout, the duodenum width of *LC^+^B^fl/fl^* mice was 1.3-fold narrower than that of the *Lgr5-Cre^+^;Bmal1^wt/wt^* (*LC^+^B^wt/wt^*) control (Figure 2B). This is due to a shorter distance between the bottom of the crypt and the top of the villi (crypt: 50.5 μM; villi: 381.3 μ M) in the *LC^+^B^fl/fl^* duodenum compared to the *LC^+^B^wt/wt^* (crypt: 55.9 μ M; villi: 441.8 μ M) (Figure 2C and 2D). Since LGR5^+^ CBCs exhibit multipotent ability to differentiate into Paneth, goblet and enteroendocrine cells (Barker et al., 2007), we evaluated the effects of *Bmal1* knockout in duodenum differentiation by examining the expression of lineage specific markers. RNA of the duodenum tissues from *LC^+^B^wt/wt^* and *LC^+^B^fl/fl^* mice were harvested at ZT5 three weeks after TAM treatment and subjected to quantitative RT-PCR (RT-qPCR) analysis (Figure S1). A significant decrease in *Lrig1* (+4 progenitor marker), *Fabp2* (enterocyte marker), and *Klf4* (goblet cell marker) expression was observed in the *LC^+^B^fl/fl^* mice. In contrast, *Muc2* level (goblet cell marker) in the duodenum was increased (Figure S1). These results demonstrated that BMAL1 contributes to the multipotent ability of LGR5^+^ CBCs and also suggested that *Bmal1* knockout in LGR5^+^ CBCs may lead to less efficient tissue regeneration and differentiation.

**Figure 2.**
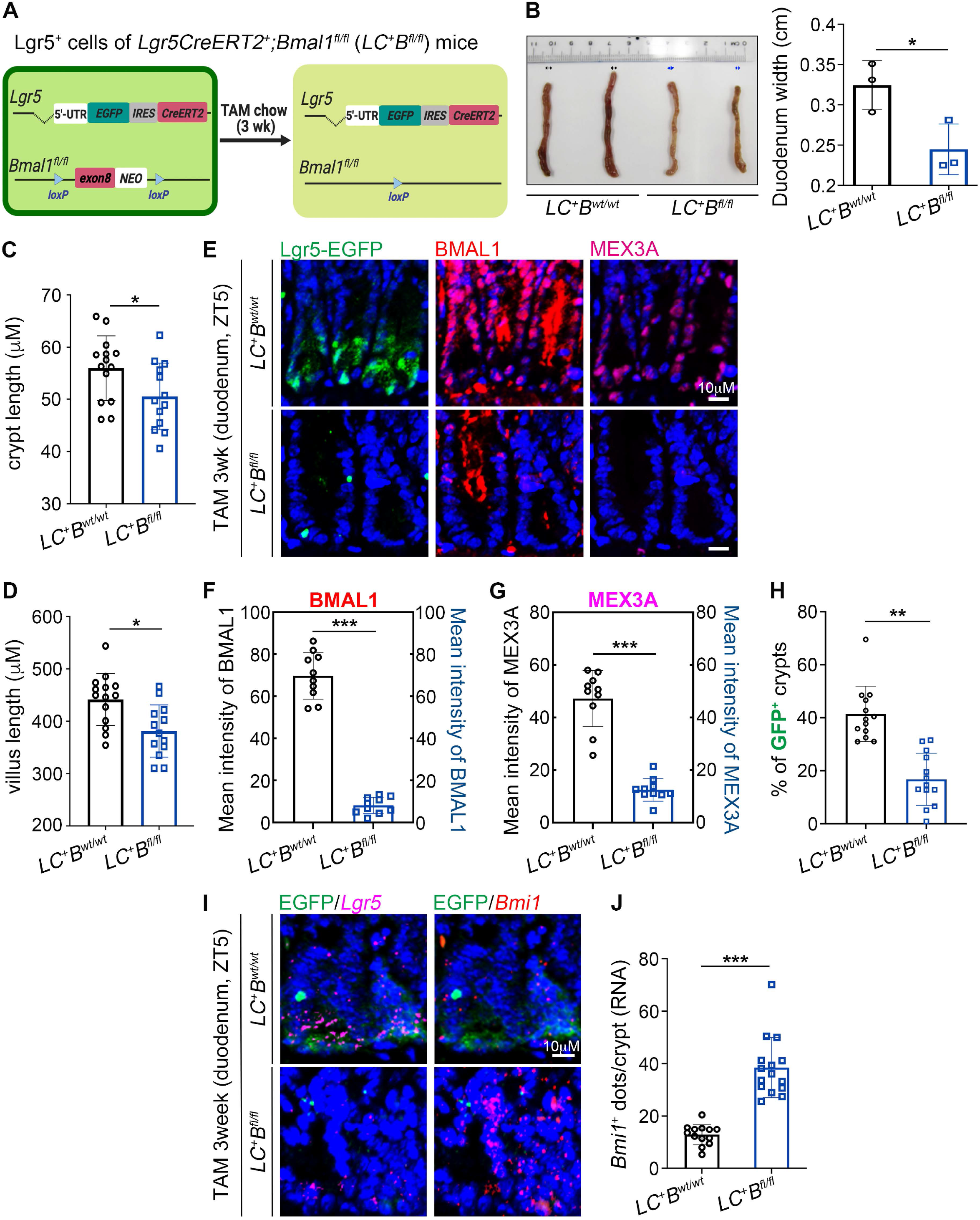
BMAL1 promotes MEX3A expression and maintains ISC homeostasis. (A) Diagram of the LGR5^+^ CBC specific BMAL1 knockout mouse model. *Lgr5CreERT2^+^;Bmal1^wt/wt^* (*LC^+^B^wt/wt^*) and *Lgr5CreERT2^+^;Bmal1^fl/fl^* (*LC^+^B^fl/fl^*) mice were fed with a tamoxifen (TAM) diet for 3 weeks to specifically knockout exon8 of the *Bmal1* gene in LGR5-EGFP^+^ CBCs. (B-D) Duodenum width (B), crypt length (C) and villus length (D) of the *LC^+^B^wt/wt^* and *LC^+^B^fl/fl^* mice. (n = 3 mice) (E-H) Immunofluorescence staining (E), quantification of BMAL1 and MEX3A expression in duodenum crypts (F, G) and prevalence of EGFP^+^ crypts (H) from TAM fed mice at ZT5. (n = 3 mice, ≥ 50 crypts/n) (I) RNAscope analysis of *Lgr5* (pink dots) and *Bmi1* (red dots) mRNA in duodenum crypts of TAM fed *LC^+^B^wt/wt^* and *LC^+^B^fl/fl^* mice. (n = 3 mice) (J) Quantification of *Bmi1* mRNA dots from (I). (n = 3 mice, ≥ 50 crypts/n) All quantification data are means ± SD, significant differences are based on unpaired *T*-tests. * *P* < 0.05; ** *P* < 0.01; *** *P* < 0.001.

In the crypts, *Bmal1* knockout led to a 3.8-fold reduction of MEX3A expression (Figure 2E-2G). Quantification of EGFP^+^ cells at the crypt base demonstrated a 2.5-fold decrease in LGR5^+^ CBC-containing crypts in the *LC^+^B^fl/fl^* mice compared to the *LC^+^B^wt/wt^* control (Figure 2E and 2H). Consistent with this result, a significant loss of *Lgr5*-RNA was observed in the crypt base of *LC^+^B^fl/fl^* mice (Figure 2I). Consistent with previous observation that BMI1^+^ +4 cells can replenish the crypt base upon loss of LGR5^+^ CBCs (Tian et al., 2011), we also found a 3-fold increase in *Bmi1* RNA in the crypts (Figure 2I, 2J). These data suggested a critical role of BMAL1 in maintaining the homeostasis between LGR5^+^ CBCs and BMI1^+^ +4 cells, likely through regulating the expression of *Lgr5* and *Mex3a*.

To further evaluate whether BMAL1 contributes to the stemness of LGR5^+^ CBCs and the succession between LGR5^+^ CBCs and BMI1^+^ +4 cells, we performed 3D-organoid culture (Sato and Clevers, 2013) using crypt cells isolated from *LC^+^B^wt/wt^* or *LC^+^B^fl/fl^* duodenum. Organoids were treated with 4-hydroxytamoxifen (4-OHT), an active metabolite of TAM, to knock out *Bmal1*. Twelve or eighteen hours after 4-OHT removal, *Bmal1*, *Mex3a*, *Lgr5* and *Bmi1* levels in the organoids were examined (Figure 3A). RT-qPCR (Figure 3B) and IF (Figure 3C and 3D) assays confirmed BMAL1 depletion in the 4-OHT-treated *LC^+^B^fl/fl^* organoids. Consistent with our observations in the TAM-treated *LC^+^B^fl/fl^* mice, EGFP^+^ (LGR5^+^) cells were significantly reduced upon *Bmal1* depletion (Figure 3C). In the *LC^+^B^fl/fl^* organoids, *Bmal11* depletion significantly decreased *Mex3a* expression compared to the control at 12 h after 4-OHT removal (Figure 3E-3G). Interestingly, the downregulated *Mex3a* level in the *LC^+^B^fl/fl^* organoids increased to similar level as in the *LC^+^B^wt/wt^* organoids 18 h after 4-OHT removal while *Bmal1* was still depleted (Figure 3E-G). Similarly, *Lgr5* level was downregulated in the *LC^+^B^fl/fl^* organoids 12 h after 4-OHT removal; however, its expression increased later (Figure 3H). BMI1^+^ +4 cells are able to proliferate and reprogram to express *Lgr5* and replenish the loss of LGR5^+^ CBCs (Tian et al., 2011). Consistent with this phenomenon, we found that *Bmi1* expression of *LC^+^B^fl/fl^* organoids was not altered 12 h after 4-OHT removal but was upregulated at the later time point (Figure 3I). This suggested that the recovery of *Lgr5* and *Mex3a* expression at this later time point was driven by reprograming of BMI^+^ cells.

**Figure 3.**
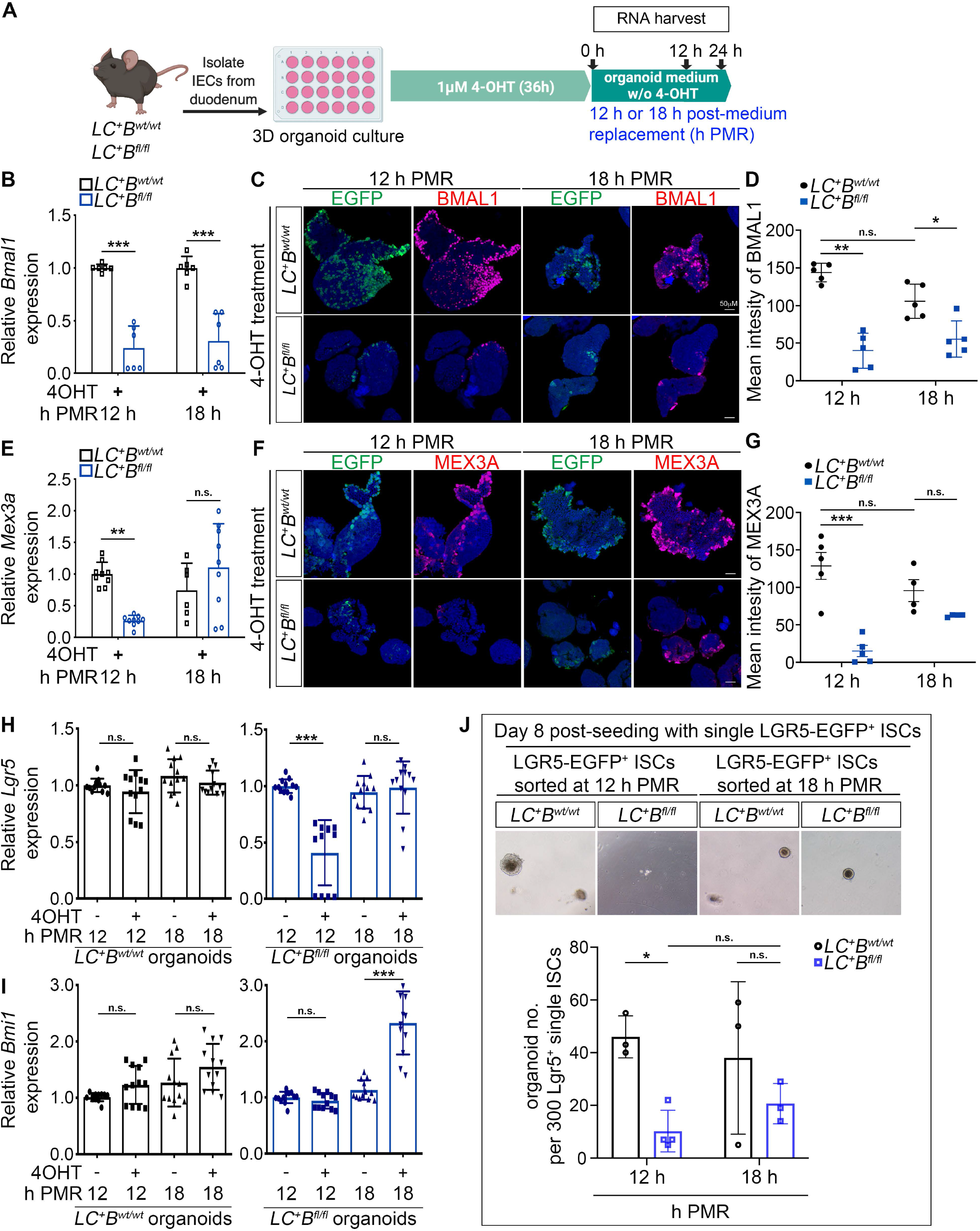
BMAL1 is essential for maintaining LGR5+-CBC self-renewal in organoids. (A) Illustration of 3D organoid culture experimental strategy using primary IECs isolated from *LC^+^B^wt/wt^* and *LC^+^B^fl/fl^* mice. 4-OHT was administrated for 36 hours. Twelve or 18 h after 4-OHT removal, organoids and RNA were harvested. Two mice were used for each group. (B) qRT-PCR analysis of *Bmal1* levels in 4-OHT treated organoids collected at 12 or 18 h post-medium replacement and 4-OHT removal (h PMR). Three independent experiments were performed and data are means ± SD from one representative experiment. (C and D) Immunofluorescence staining (C) and quantification of BMAL1 (D) protein level in organoids from 12 or 18 h PMR samples. EGFP^+^ cells are LGR5-expressing CBCs. (n ≥ 7 organoids per time point). (E) qRT-PCR analysis of *Mex3a* levels in 4-OHT treated organoids collected at 12 or 18 h PMR. Three independent experiments were performed and data are means ± SD from one representative experiment. (F and G) Immunofluorescence staining (F) and quantification of MEX3A (G) protein level in organoids from 12 or 18 h PMR samples. (n ≥ 7 organoids per time point). (H and I) qRT-PCR analysis of *Lgr5* (H) and *Bmi1* (I) expression in 4-OHT treated (4OHT +) or non-treated (4OHT-) organoids collected at 12 or 18 h PMR (4OHT removal for the 4OHT+ treatment). Three independent experiments were performed and data are means ± SD from one representative experiment. (J) The number of secondary organoids formed after isolation of LGR5-EGFP^+^ CBCs from 4-OHT treated organoids at 12 or 18 h PMR and culture for 8 days. Three hundred sorted LGR5-EGFP^+^ CBCs were seeded per well in growth factor reduced matrigel. All quantification data are means ± SD, significant differences are based on two-way ANOVA. * *P* < 0.05; ** *P* < 0.01; *** *P* < 0.001; n.s., no statistically significant difference.

We further evaluated the role of BMAL1 in self-renewal ability of LGR5^+^ CBCs by performing a secondary 3D-organoid forming assay using EGFP^+^ cells isolated from the organoids at 12 (early) or 18 (late) h after 4-OHT removal. For the EGFP^+^ cells isolated from the *LC^+^B^wt/wt^* organoids at both time points, 13-15 percent successfully formed secondary organoids by eight days after isolation (Figure 3J). In contrast, the organoid forming efficiency of *Bmal1*-depleted EGFP^+^ cells isolated at the 12 h time point was significantly impaired (only 10.3 out of 300 cells formed secondary organoids). Importantly, at the later 18 h time point, the *Bmal1*-depleted EGFP^+^ cells with elevated *Bmi1* expression (Figure 3I) had an increased trend of organoid forming ability (Figure 3J, *p* = 0.8). These results, together with the observations from the TAM-treated *LC^+^B^fl/fl^* mice, indicated that immediately after loss of BMAL1, *Mex3a* and *Lgr5* expression were both downregulated and the prevalence of LGR5^+^ CBCs decreased. Thus, BMAL1 expression was critical to promote LGR5 expression and maintain the self-renewal activities of LGR5^+^ CBCs. At later times, BMI1^+^ +4 cells may proliferate and reprogram to express *Lgr5*, possibly via upregulated *Mex3a* expression, to compensate for the loss of LGR5^+^ CBCs.

### BMAL1 and MEX3A co-regulate LGR5 expression in intestinal epithelial cells

The results described above demonstrated that *Bmal1* knockout led to decreased MEX3A and LGR5 levels *in vivo*. To further test whether BMAL1 regulated *Mex3a* and *Lgr5* expression, *Bmal1* expression was depleted in an immortalized mouse intestinal epithelial cell line (mIEC) using lentiviral shRNA transduction. A decrease in MEX3A and LGR5 but a mild increase in BMI1 protein levels were observed upon *Bmal1* knockdown (Figure 4A). Consistent with the observation that *Mex3a* knockout decreased LGR5^+^ CBCs (Pereira et al., 2020), depletion of *Mex3a* with shRNA also led to a decrease in LGR5 protein expression and a slight increase in BMI1 (Figure 4B). These data indicated that both BMAL1 and MEX3A contribute to LGR5 expression and may either directly or indirectly suppress BMI1 expression.

**Figure 4.**
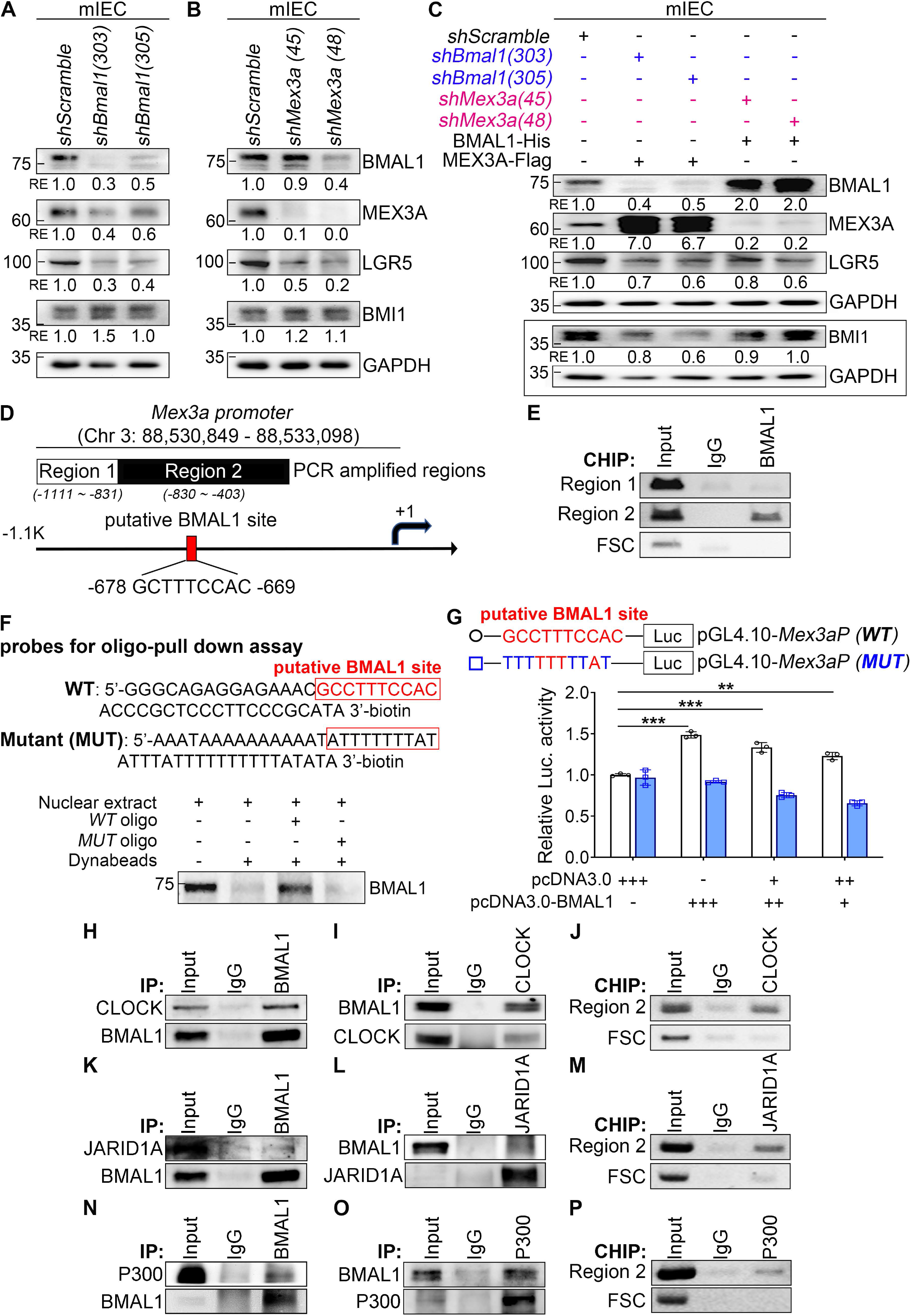
BMAL1 exerts an opposite regulatory effect on LGR5 and BMI1 levels in mIEC by transcriptionally upregulating *Mex3a*. (A and B) Immunoblot of BMAL1, MEX3A, LGR5 and BMI1 in mIEC transduced with *shBmal1* (A) or *shMex3a* (B) lentivirus. *shScramble* was used as a transduction control and GAPDH was used as an internal control and blot loading control. Blots shown are from one representative experiment of two independent experiments. Numbers below each blot are the quantification of band intensity relative to the *shScramble* control. (C) Immunoblot of BMAL1, MEX3A, LGR5 and BMI1 in *shBmal1* or *shMex3a* transduced IECs with MEX3A or BMAL1 overexpression, respectively. *shScramble* was used as a transduction and RNAi control and GAPDH was used as an internal control. Blots shown are from one representative experiment of two independent experiments. Numbers below each blot are the quantification of band intensity relative to the *shScramble* control. (D) Diagram shows one predicted BMAL1 binding site (red box) on the region 2 (−830 to −403) of the mouse *Mex3a* promoter using the JASPAR database. (E) ChIP analysis of BMAL1 occupancy on the *Mex3a* promoter in mIEC. Mouse IgG and far site control region (FSC; Chr 3, 88,521,891-88,522,294) were used as negative controls. Gels shown are from one representative experiment of two independent experiment. (F) Biotinylated oligo pull-down of BMAL1 using WT or mutant predicted BMAL1 binding sequence and mIEC nuclear extract. Blots shown are from one representative experiment of three independent experiments. (G) Top: Schematic diagram of pGL4.10-*Mex3a* promoter-Luc reporter with wild-type BMAL1 binding sequences. Bottom: Luciferase reporter assays were conducted using 293T cells co-transfected with pcDNA3.0-BMAL1 and the wild type (WT, black bars) or mutant (MUT, blue bars) BMAL1 promoter constructs. Three replicate experiments were performed. Data show means ± SD based on two-way ANOVA. ** *P* < 0.01, *** *P* < 0.001. (H-J) Co-IP (H), reciprocal-IP (I) and ChIP (J) analysis of BMAL1 and CLOCK on the *Mex3a* promoter using nuclear extract from mIEC. (K-M) Co-IP (K), reciprocal-IP (L) and ChIP (M) analysis of BMAL1 and JARID1A on the *Mex3a* promoter using nuclear extract from mIEC. (N-P) Co-IP (N), reciprocal-IP (O) and ChIP (P) analysis of BMAL1 and P300 on the *Mex3a* promoter using nuclear extract from mIEC. For (H-P), IgG and far site region (FSC) were used as an IP control. Blots and gels shown are from one representative experiment of three independent experiments.

To further evaluate whether MEX3A is a critical factor of BMAL1-mediated LGR5 upregulation, pBMAL1-His or pMEX3A-Flag plasmid was transiently transfected into *Mex3a-* or *Bmal1-*depleted mIEC, respectively. Overexpressing either BMAL1 in *Mex3a*-depleted cells or MEX3A in *Bmal1*-depleted cells partially compensated the reduced LGR5 resulted from *Bmal1* or *Mex3a* knockdown (Figure 4C). In addition, the increase in BMI1 expression observed in the *Bmal1* or *Mex3a* knockdown cells was abolished when MEX3A or BMAL1 was overexpressed, respectively (Figure 4C). Together, these results suggested that BMAL1 and MEX3A co-regulate LGR5 and BMI1 expression in intestinal crypts.

### BMAL1 directly upregulates MEX3A transcription

Given the close relationship between *Bmal1* depletion and MEX3A downregulation (Figure 4A and 4B), we analyzed the *Mex3a* promoter using the JASPAR database (Fornes et al., 2020) and found a putative BMAL1 binding site [GCTTTCCAC; −678~-669 nt upstream of the transcriptional start site (TSS)] (Figure 4D). Even though this site is not a complete match to the BMAL1 consensus binding site [E-box (E1, CACGTG) or E1-E2 (E2, AACGTG) tandem sites] (Rey et al., 2011), chromatin-immunoprecipitation (ChIP) assay found BMAL1 associated with this *Mex3a* promoter region (Figure 4E). Biotinylated-oligonucleotide-pull-down assay demonstrated that BMAL1 directly and specifically bound to this site, as BMAL1 bound an oligo probe containing this putative site but not a mutated probe (Figure 4F). Transient reporter assay showed that mutation of this site abolished BMAL1 transactivation of the *Mex3a* promoter in HEK-293T cells (Figure 4G). In addition, Co-IP and ChIP assays found that BMAL1 interacted with the core circadian regulator CLOCK (Figure 4H-4J), histone demethylase JARID1A (Figure 4K-4M) and co-activator P300 (Figure 4N-4P) on the *Mex3a* promoter to activate *Mex3a* expression. Together, these results indicated that BMAL1 directly activates *Mex3a* expression.

### BMAL1 directly activates *Lgr5* expression and indirectly stabilizes *Lgr5* mRNA via MEX3A upregulation

Since depletion of *Bmal1* decreased LGR5 level in mIEC (Figure 4A), we also investigated whether BMAL1 directly regulated LGR5 expression by binding to the *Lgr5* promoter. Promoter analysis using the JASPAR database identified 4 putative BMAL1 binding sites located between - 2020 nt and −694 nt upstream of the TSS of the *Lgr5* promoter (Figure 5A). BMAL1 was associated with two regions (region 3 and 4) containing three predicted BMAL1 sites (Figure 5B). Similar to the *Mex3a* promoter, CLOCK (Figure 5C) and JARID1A (Figure 5D), but not P300 (Figure 5E), were found to co-localize with BMAL1 on region 4 of the *Lgr5* promoter. RT-qPCR further confirmed that *Bmal1* depletion resulted in downregulation of *Lgr5* gene expression (Figure 5F). Together, these data indicated that BMAL1 transcriptionally regulates *Lgr5* expression in the crypts.

**Figure 5.**
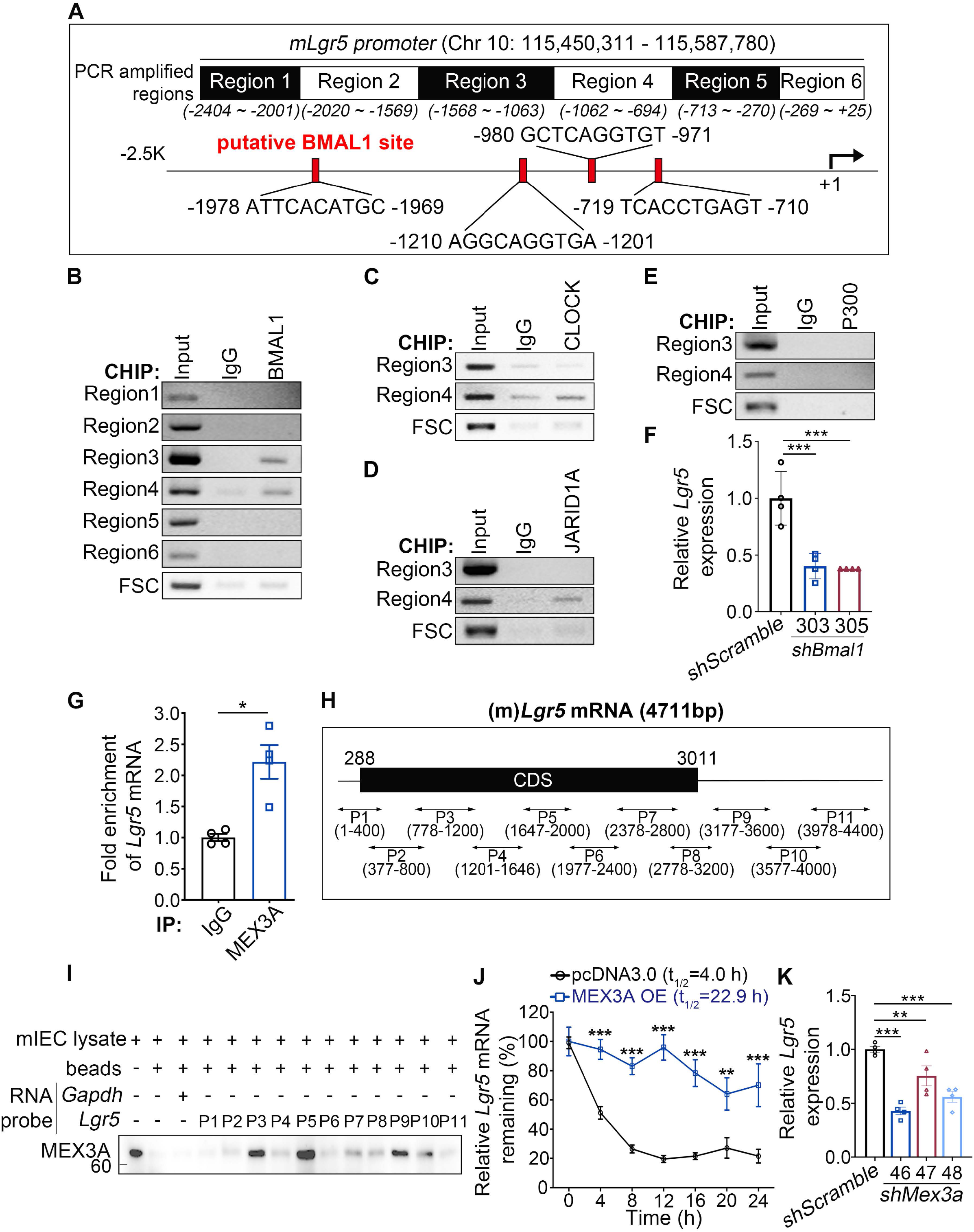
BMAL1 directly binds the *Lgr5* promoter to drive its expression and upregulate *Mex3a* to stabilize *Lgr5* mRNA. (A) Four putative BMAL1 binding sites (red boxes) were found in region 2 to region 4 (−2020 to −694) of the mouse *Lgr5* promoter. (B-E) ChIP analysis of BMAL1 (B), CLOCK (C), JARID1A (D) and P300 (E) occupancy on the *Lgr5* promoter in mIEC. IgG and far site region (FSC; Chr 10, 115,443,315-115,443,757) were used as a control for the ChIP experiments. Gels shown are from one representative experiment of two independent experiments. (F) qRT-PCR of *Lgr5* in mIEC transduced with *shScramble* control or *shBmal1* lentiviruses (clone #303 or #305). Three independent experiments were performed. Data show means ± SD based on one-way ANOVA. *** *P* < 0.001. (G) RNA-immunoprecipitation assay of MEX3A-*Lgr5* complex using mIEC lysates. IgG was used as an antibody control for immunoprecipitation. Three independent experiments were performed. Data are presented as means ± SD, significant differences are based on Student’s t test. * *P* < 0.05. (H) Schematic of mouse *Lgr5* mRNA and eleven biotinylated RNA probes (P1 to P11) spanning nearly whole *Lgr5* mRNA sequence used for RNA pull-down assay. (I) RNA pull-down assay of MEX3A using 11 biotinylated RNA probes that covered the *Lgr5* coding regions (CDS) and 3’-UTR. Beads only and the *Gapdh* RNA probe were used as pull-down controls. Blot shown is from one representative experiment of three independent experiments. (J) 5-bromouridine (BrU) immunoprecipitation chase assay (BRIC) of *Lgr5* mRNA in mIEC transfected with pcDNA3.0 control vector or pMex3a-3X flag. *Lgr5* levels at 0 h was set as 100%. Two independent experiments were performed. Data show means ± SD, significant differences are based on two-way ANOVA. ** *P* < 0.01, *** *P* < 0.001. (K) qRT-PCR analysis of *Lgr5* levels in mIEC transduced with *shScramble* or *shMex3a* lentiviruses (clone #46, #47 or #48). Three independent experiments were performed. Data show means ± SD, significant differences are based on one-way ANOVA. ** *P* < 0.01, *** *P* < 0.001.

In the BMAL1 expressing mIEC, *Mex3a* depletion also reduced LGR5 level (Figure 4B). RNA immunoprecipitation (RNA-IP) assay showed that MEX3A protein bound to *Lgr5* mRNA (Figure 5G). This suggested that MEX3A may regulate *Lgr5* expression post-transcriptionally. Since the structure and RNA binding motif of MEX3A remain unknown, we used eleven biotinylated-RNA probes (400-nt to 500-nt in length per probe) spanning the *Lgr5* mRNA including the 3’-UTR (Figure 5H) to perform biotinylated-RNA pull-down assay. The results showed that MEX3A interacted with *Lgr5* mRNA at multiple sites, including two in the coding region (CDS, probes 3, 5) and one in the 3’-UTR (probe 9) (Figure 5I). MEX3A binding stabilized *Lgr5* mRNA as MEX3A overexpression significantly increased the half-life of *Lgr5* mRNA from 4 to 22.9 hours in mIEC (Figure 5J). Consistent with these observations, depletion of MEX3A using lenti-viral transduced shRNAs reduced *Lgr5* mRNA level in mIEC (Figure 5K). Together, these data indicated that BMAL1 could directly activate *Lgr5* transcription and indirectly stabilize *Lgr5* mRNA by upregulating MEX3A.

### BMAL1 and MEX3A contribute to *Bmi1* suppression

Since increased BMI1 level was observed in the *Bmal1* or *Mex3a* knockdown mIEC (Figure 4A, 4B), BMAL1 and MEX3A may also maintain LGR5^+^ CBCs by inhibiting BMI1 expression. Two E-boxes and two putative BMAL1 binding sites were identified between −980 nt and +26 nt of the *Bmi1* promoter (Figure S2A). BMAL1 was associated with the site closest to the −1 kb region (region 5) (Figure S2B). BMAL1 binding was associated with *Bmi1* suppression as *Bmal1* depletion led to elevated *Bmi1* mRNA level (Figure S2C).

Also, RNA-IP and RNA pull-down assay showed that *Bmi1* mRNA was bound by MEX3A at both the CDS (region 1) and 3’-UTR (region 5) regions (Figure S2D-S2F). However, unlike the effect of MEX3A binding to *Lgr5* mRNA, overexpression of MEX3A did not alter *Bmi1* mRNA stability (Figure S2G). Depletion of *Mex3a* also did not affect *Bmi1* mRNA level (Figure S2H). Importantly, BMI1 protein was more stable in *Mex3a* depleted mIEC after cycloheximide (CHX) treatment compared to the control (Figure S2I). These data suggested that MEX3A contributed to BMI1 suppression at the protein level, rather than the post-transcriptional level, probably though its RING domain at the c-terminus which is involved in protein degradation (Bufalieri et al., 2020; Cano et al., 2010).

### BMAL1 and MEX3A expression reprogram BMI1+ +4 cells to express LGR5

It is well established that BMI1^+^ +4 cells can convert into LGR5-expressing cells upon intestinal damage (Tian et al., 2011; Yan et al., 2012). Since BMAL1 and MEX3A are critical for LGR5 expression and LGR5^+^ CBC maintenance (Figure 2, 3), it is likely that BMAL1 and MEX3A contribute to *Lgr5* expression during BMI1^+^ +4 cell reprogramming. To test this hypothesis, we established a BMI1^+^ lineage tracing mIEC line (Bmi1-P2A-cherry mIEC). A P2A-mCherry reporter was fused to the end of the exon 10 of *Bmi1* gene using a CRISPR-Cpf1 system (Figure S3A) and the mCherry^+^ mIEC cells were confirmed to also express BMI1 (Figure S3B). The prevalence of LGR5^+^cells among the mIEC was 0.9% (± 0.36) (Figure 6A). Note that we found two discrete subpopulations of BMI1^+^ (mCherry^+^) mIEC cells, Bmi1-mCherry^High^ (3.9% ± 1.16) and Bmi1-mCherry^Low^ (13% ± 2.21) (Figure 6B). To investigate whether BMAL1 and MEX3A were differentially expressed between LGR5^+^ and BMI1^+^ subpopulations, immunoblot analysis was performed using LGR5^+^ and Bmi1-mCherry^High^ cells with 71.9% ± 12.54 and 91.6% ± 4.65 purity, respectively (Figure 6A and 6B). As expected, higher expression of BMAL1 and MEX3A were found in the LGR5^+^ cells compared to the Bmi1-mCherry^High^ cells (Figure 6C). These results suggested that BMAL1 and MEX3A may be required to drive the reprogramming of BMI1^+^ +4 cells to express LGR5.

**Figure 6.**
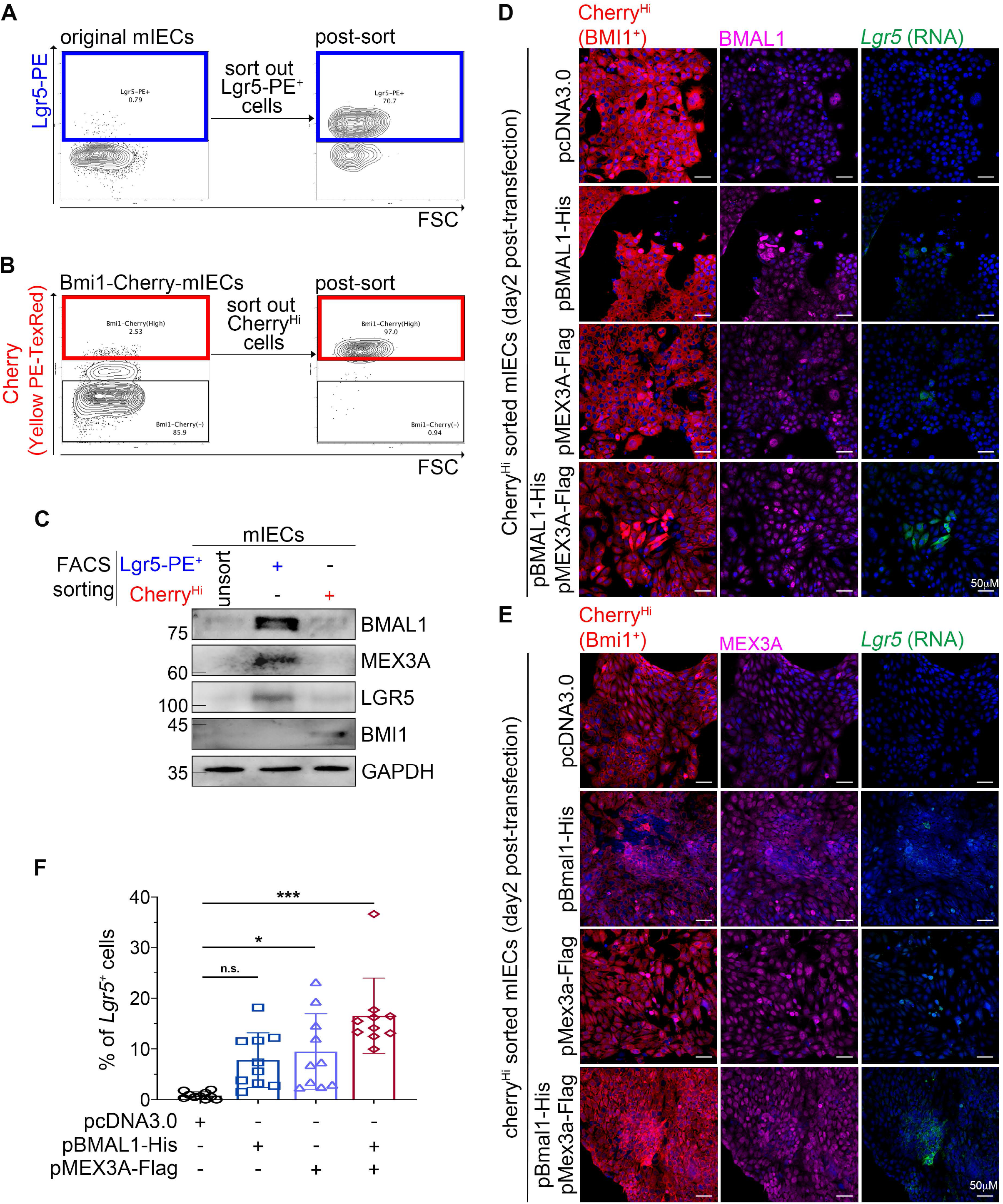
BMAL1 and MEX3A promote reprogramming of BMI1+ cells to express LGR5. (A and B) Enrichment of LGR5^+^ (PE^+^) or Bmi1-expressing^High^ (Cherry^High^) ISCs from parental or CRISPR Bmi1-P2A-mCherry knock-in mIEC (Bmi1-Cherry-mIEC) using FACS sorting. The contour plots showed the percentage of LGR5-PE^+^ (blue frame) and Bmi1-Cherry^High^ (red frame) subpopulations before and after sorting. (C) Immunoblot of BMAL1 and MEX3A protein levels in sorted LGR5-PE^+^ and Bmi1-Cherry^High^ subpopulations. Unsorted mIEC were used as a control for detecting the basal level of BMAL1 and MEX3A. GAPDH was used as a loading control. Blots shown are from one representative experiment of at least two independent experiments. (D and E) Immunofluorescence staining of BMAL1 (D) and MEX3A (E) in sorted Bmi1-Cherry^High^ subpopulation transduced with pBMAL1-His, pMEX3A-Flag, or both. *Lgr5* expression was detected by RNAscope analysis. pcDNA3.0 was used as a transfection control. Three independent experiments were performed. (F) Quantification of LGR5^+^ cells in Bmi1-Cherry^High^ subpopulation ectopically expressing BMAL1, MEX3A or both from (D) and (E). Data show means ± SD, significant differences are based on one-way ANOVA. * *P* < 0.05, *** *P* < 0.001, n.s., no statistically significant difference.

To further evaluate if BMAL1 and MEX3A promote BMI1^+^ +4 cells reprogramming to LGR5 expressing cells, BMI1-mCherry^High^ cells were sorted and transfected with either pBmal1-His alone, pMex3a-Flag alone or both plasmids for two days. Increased fluorescence signals of BMAL1 (Figure 6D) and MEX3A (Figure 6E) confirmed the ectopic expression of these two proteins in the BMI1-mCherry^High^ cells. Upon BMAL1 or MEX3A ectopic expression, approximately 7.8% or 9.4% of the BMI1-mCherry^High^ cells converted to *Lgr5*-expressing cells, respectively. Co-expression of BMAL1 and MEX3A further increased the prevalence of the *Lgr5*-expressing cells to 16.5% (Figure 6D-6F). These results demonstrated that BMAL1 and MEX3A can reprogram BMI1-mCherry^High^ cells to express LGR5. Together, our findings indicated an important role of BMAL1-MEX3A co-regulation in maintaining the homeostasis and succession between LGR5^+^ CBCs and BMI1^+^ +4 cells.

### 5-FU delivery when *Bmi1* reached its peak level in crypt cells protected ISCs from apoptosis

In the crypt cells, rhythmic expression of *Lgr5* and *Bmi1* were co-regulated by BMAL1 and MEX3A (Figure 1). *Lgr5* reached the peak level at ZT9 during the subjective day (Figure 1E) and dropped to the low level at ZT17 during the subjective night when *Bmi1* reached its peak level (Figure 1F). Also, BMI1 has been identified as an early DNA damage response protein (Ismail et al., 2010) and a factor promoting resistance to radiation (Facchino et al., 2010). Such protein function may explain why BMI1^+^ +4 cells are more resistant to intestinal injury (Keefe et al., 2000; Yan et al., 2012). Our analysis suggested that, 5-FU delivery at the time when *Bmi1* reaches the peak level at subjective night may result in less chemotherapy-induced side effects. To explore this possibility, *Lgr5-Cre* mice kept under LD condition were separated randomly into two groups and subjected to 0.4mg/kg 5-FU treatment (a mouse equivalent dose extrapolated from human based on body surface area) (Nair and Jacob, 2016) by jugular catheter injection every other day for four weeks. 5-FU was delivered at either ZT9 or at ZT17. After four weeks of consecutive treatment, all mice were sacrificed at ZT5 and the stress response of crypt cells was evaluated (Figure 7A). Compared to the PBS-treated control, 5-FU treatment significantly reduced proliferation of LGR5^+^ CBCs in both groups (Figure 7B, 7C). However, mice treated with 5-FU at ZT17 maintained a higher prevalence of proliferating LGR5^+^ CBCs (22%) than mice treated at ZT9 (3.8%; Figure 7C, compare blue square to black circle). Even more striking was the observation that 5-FU-induced apoptosis could only be detected in the mice treated at ZT9. In mice treated at ZT9, approximately 17.8% of the villus cells and 3.3% of the crypt cells were apoptotic (Figure 7D-7F, black circle). In contrast, no significant increase in apoptotic cells was observed in the ZT17 group and only 1.8% of the villus cells and 0.2% of the crypt cells were TUNEL positive (Figure 7D-7F, blue square). These results indicated that delivery of 5-FU when *Lgr5* level is low and *Bmi1* expression is high could protect normal intestinal epithelium and ISCs from chemotherapy-induced damage.

**Figure 7.**
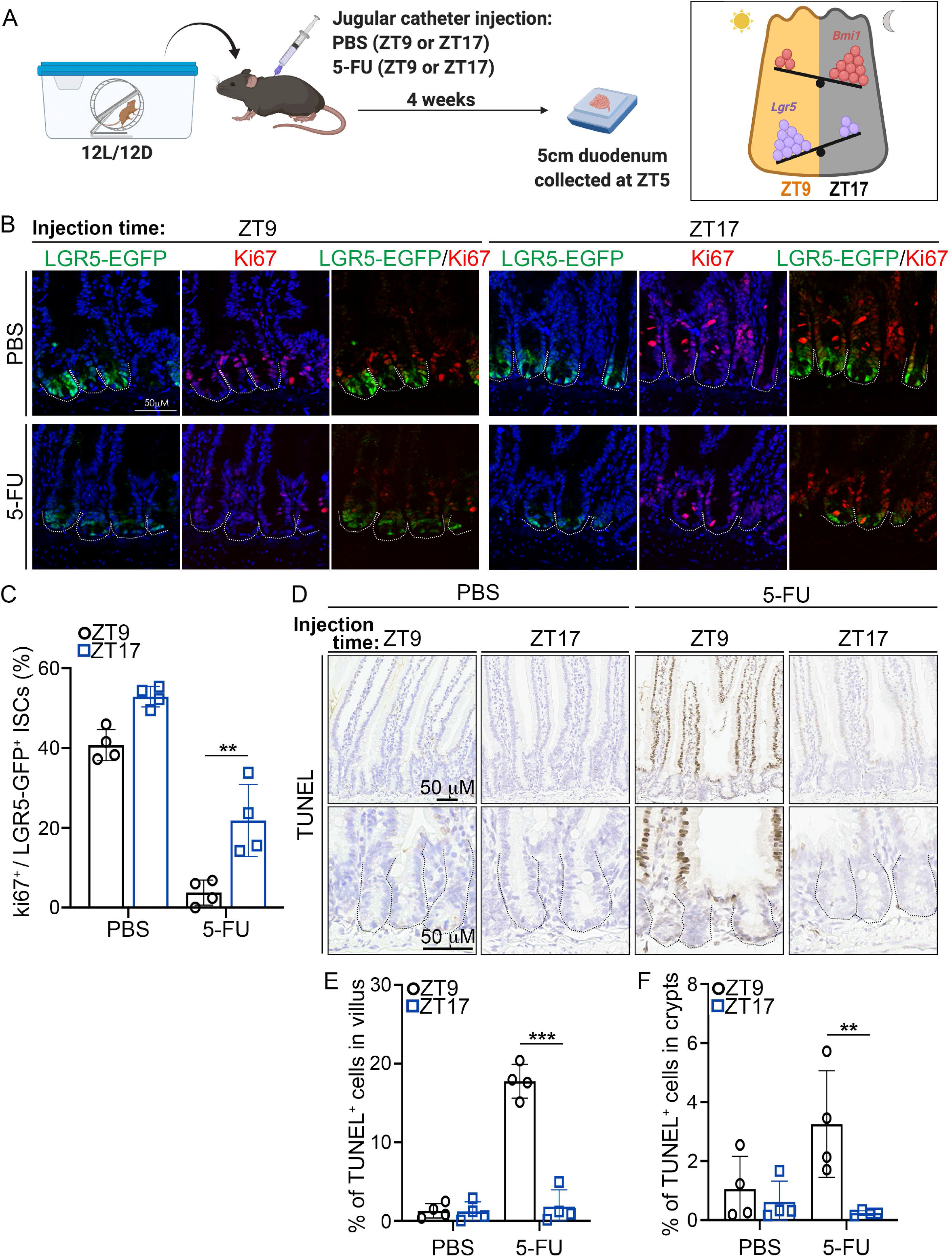
5-FU delivery at ZT17 protects crypt cells from apoptosis and allows sustained ISC cell proliferation. (A) Diagram of the chrono-modulated 5-FU jugular catheter injection model. *Lgr5*-*Cre*^+^ mice were implanted with catheters to the external jugular vein and kept under LD condition. 5-FU was injected at ZT9 when *Lgr5* reached the peak level, or ZT17 when *Bmi1* was high (inset) every other day. Mice were sacrificed at ZT5, 4 weeks after the 5-FU or PBS control treatments. (B) Immunofluorescence staining of a proliferation marker (Ki67) and EGFP (LGR5^+^ ISCs) in duodenum from PBS or 5-FU treated *Lgr5*-Cre^+^ mice. Representative images are shown, two independent experiments were performed. (C) Quantification of Ki67^+^ percentages in LGR5-EGFP^+^ CBCs in (B). Data show means ± SD based on one-way ANOVA. ** *P* < 0.01 (n = 2 mice/treatment; ≥ 55 crypts/n) (D) TUNEL staining assay of apoptotic cells in duodenum from PBS or 5-FU treated *Lgr5*-Cre^+^ mice. (n= 2) (E-F) Quantification of TUNEL^+^ cells in all cells counted in villus (E) and crypts (F). Data shown means ± SD, significant differences are based on one-way ANOVA. ** *P* < 0.01, *** *P* < 0.001 (n = 2 mice/treatment; ≥ 55 crypts/n)

Emerging evidence has indicated that the circadian clock modulates intestinal physiology, such as IEC regeneration (Karpowicz et al., 2013), microbiota-IEC crosstalk (Mukherji et al., 2013), and intestinal permeability (Summa et al., 2013). BMAL1 is essential for maintaining rhythmic cell division within intestinal organoids (Matsu-Ura et al., 2016). However, it has been unclear whether BMAL1 regulates the succession or regeneration between CBCs and +4 ISCs. Using LGR5^+^ CBC specific *Bmal1* knockout mouse, organoid and mIEC model, we showed that co-regulation of BMAL1 and MEX3A is crucial to maintain homeostasis between the fast-proliferating LGR5^+^ CBCs and damage-resistant BMI1^+^ +4 cells. Ablation of *Bmal1* reduced both LGR5^+^ CBCs and MEX3A expression but increased BMI1 levels. Induction of BMAL1 and MEX3A was sufficient to promote reprogramming of BMI1^+^ cells to express *Lgr5* (Figure S4). Moreover, delivery of 5-FU at ZT17 when *Bmi1* reached the peak level in the crypts protected ISCs from apoptosis. These results uncovered a significant role of BMAL1 in ISCs to regulate ISC homeostasis and regeneration. The data also provided a biological basis for developing a 5-FU chronotherapeutic strategy to mitigate apoptotic side-effects in intestinal cells.

Consistent with previous observations that MEX3A is essential for the survival of LGR5^+^ cells and organoid forming abilities (Pereira et al., 2020), our study further demonstrated that MEX3A is transcriptionally upregulated by BMAL1 (Figure 4) and plays an essential role to coregulate ISC homeostasis with BMAL1 (Figure 2, 3). MEX3A is a unique protein with dual functions since it is not only a RBP that binds and regulates RNA stability via its KH1-KH2 domains (Pereira et al., 2013), but also a E3 ubiquitin ligase that promotes protein poly-ubiquitination via its RING domain at the c-terminus (Bufalieri et al., 2020; Cano et al., 2010). Our data suggested that MEX3A utilized both functions to regulate the homeostasis of ISCs and regeneration in the crypts. MEX3A bound the *Lgr5* mRNA to increase its stability (Figure 5). Furthermore, its expression contributed to BMI1 protein destabilization (Figure S2). Since *Bmal1* knockout in LGR5^+^ CBCs perturbed expression of many lineage specific differentiated markers (Figure S1), whether MEX3A also participates in ISC differentiation and regulates lineage specific IEC functions are worthy topics for further investigation.

Cancer chronotherapy is a novel therapeutic strategy to deliver chemotherapeutic agents at specific circadian time points to maximize drug efficacy and minimize adverse effects (Levi et al., 2010). Several random trials have tested chrono-modulated combination treatment of 5-FU, oxaliplatin and leucovorin/folinic acid and found that drug delivery between 2 and 4 o’clock in the morning, instead of constant infusion, showed higher tumor objective response rates (>50%) (Levi et al., 1997; Mormont and Levi, 2003) and milder mucositis (Levi et al., 1997) in metastatic colon cancer patients. The time-dependent differences in treatment efficacy could be due to fewer S-phase cells (Smaaland et al., 1992; Smaaland et al., 1991) and increased activity of dehydropyrimidine dehydrogenase, a rate-limiting enzyme for 5-FU catabolism (Harris et al., 1990). Our results provide additional mechanisms underlying the chrono-response to 5-FU. Modulation of ISC homeostasis and regulation LGR5 and BMI1 expression by BMAL1 and MEX3A may contribute to protection against apoptosis in response to 5-FU treatment. This observation is in line with previous studies indicating that BMAL1 and MEX3A, as well as BMI1, play roles in cell division and damage-induced repair and regeneration.

In intestinal crypt cells, BMAL1 maintains rhythmic cell proliferation under unstressed condition and facilitates regeneration upon radiation injury (Stokes et al., 2017). In mammary epithelial cells, BMAL1 depletion enhanced DNA damage-induced apoptosis (Korkmaz et al., 2018). Consistent with these phenomena, we found that 5-FU-induced apoptosis was most severe at ZT9 when *Lgr5* expression was high while BMAL1 reached the trough level (Figure 1E, 7D-7F). Moreover, LGR5^+^ CBCs can be divided into two subpopulations with different proliferation rate and stress response based on MEX3A expression level (Barriga et al., 2017). The MEX3A^high^-LGR5^+^ cells are more stress resistant and critical for intestinal epithelium repair and homeostasis upon toxic insults (Barriga et al., 2017). The stress resistant property of the MEX3A^high^-LGR5^+^ cells is consistent with our data showing that 5-FU delivery at ZT17 was less detrimental when both *Mex3a* and BMAL1 expression were high (Figure 1D, 7D-7F). Furthermore, BMI1 is known to promote cell proliferation (Jacobs et al., 1999), facilitate stem cell self-renewal (Yadirgi et al., 2011) and initiate DNA repair upon damage (Ismail et al., 2010). This also explains why 5-FU delivery at ZT17 when *Bmi1* expression was high protected ISCs from apoptosis (Figure 1F, 7D-7F). Together, high BMAL1, MEX3A and BMI1 expression at ZT17 may set up the stage for immediate damage repair and therefore protect cells from apoptosis in response to 5-FU treatment.

It has noted that more than 200 transcripts cycle with an ultradian period (~12 h) in mouse livers (Hughes et al., 2009). Consistent with this notion, we observed that BMAL1, *Mex3a* and *Lgr5* displayed an ultradian rhythm in intestinal crypt cells, whereas *Bmi1* transcripts showed a 24 h oscillation. A recent study further confirmed that the ultradian genes in mouse liver are enriched in the endoplasmic reticulum (ER) and mitochondria metabolism pathways (Zhu et al., 2017), suggesting the peripheral 12 h clock might modulate the ER and mitochondrial functions. In addition, it has been shown that intestinal LGR5^+^ CBCs have higher mitochondrial activity which affects their organoid forming abilities (Rodriguez-Colman et al., 2017). Additionally, an organoid culture system also showed rhythmic crypt budding events with a period of 12 h (Matsu-Ura et al., 2016), which give rise to the hypothesis that the ultradian oscillation in crypt cells might contribute to stem cell budding and crypt formation. As ultradian gene expression is associated with mitochondria metabolic activity and mitochondria activities are critical for ISC functions, we speculate that the ultradian gene expression in intestinal crypt stem cells might play essential roles in regulating their function and should be further explored.

Overall, our study demonstrates how the circadian core transcriptional factor, BMAL1, and MEX3A co-regulate the succession between the intestinal LGR5^+^ and BMI1^+^ stem cells and contribute to maintaining self-renewal activities of LGR5^+^ CBCs. Importantly, treatment of 5-FU at ZT17 when BMAL1 and *Mex3a* reach high level protects ISCs against apoptosis and permits ISCs to continue proliferating. This indicated that BMAL1 and MEX3A are essential for sustaining ISC survival following toxic insults. These findings not only demonstrate the important function of BMAL1 in stem cell biology, but also provide a basis for developing chronotherapeutic strategy to mitigate chemotherapeutic side effects.

### Limitations of Study

We showed that BMAL1 played a key role in maintaining LGR5^+^ CBCs and participated in regulating the succession between LGR5^+^ CBCs and BMI1^+^ +4 cells via upregulating *Mex3a* and *Lgr5* expression. Our study focused on the succession between LGR5^+^ CBCs and BMI1^+^ +4 cells. We did not include other cell populations which can replenish the crypts upon ablation of LGR5^+^ CBCs. As new evidence has suggested that LGR5^+^ CBCs can also be derived from dedifferentiation of absorptive and secretory progenitors, enterocyte precursors or Paneth cells (Murata et al., 2020; Schmitt et al., 2018; Tetteh et al., 2016; Yu et al., 2018) in response to damage, it should be determined whether BMAL1 also involved in these dedifferentiation process. In addition, the differential mechanisms underlying BMAL1 ultradian oscillation in the crypts and circadian rhythmic expression in the villus should be further investigated. As a proof of concept, we administered 5-FU at ZT9 when *Lgr5* was high or at ZT17 when *Bmi1* reached high level to examine the stress response in normal intestinal cells. Future studies will need to utilize orthotopic xenograft colon cancer models to test the cancer killing efficacy and side effects of 5-FU at different time points in addition to ZT9 and ZT17 to develop a mechanism-based chrono-chemotherapy regimen.

## STAR★METHODS

Detailed methods are provided in the online version of this paper and include the following:

### Key Resource Table

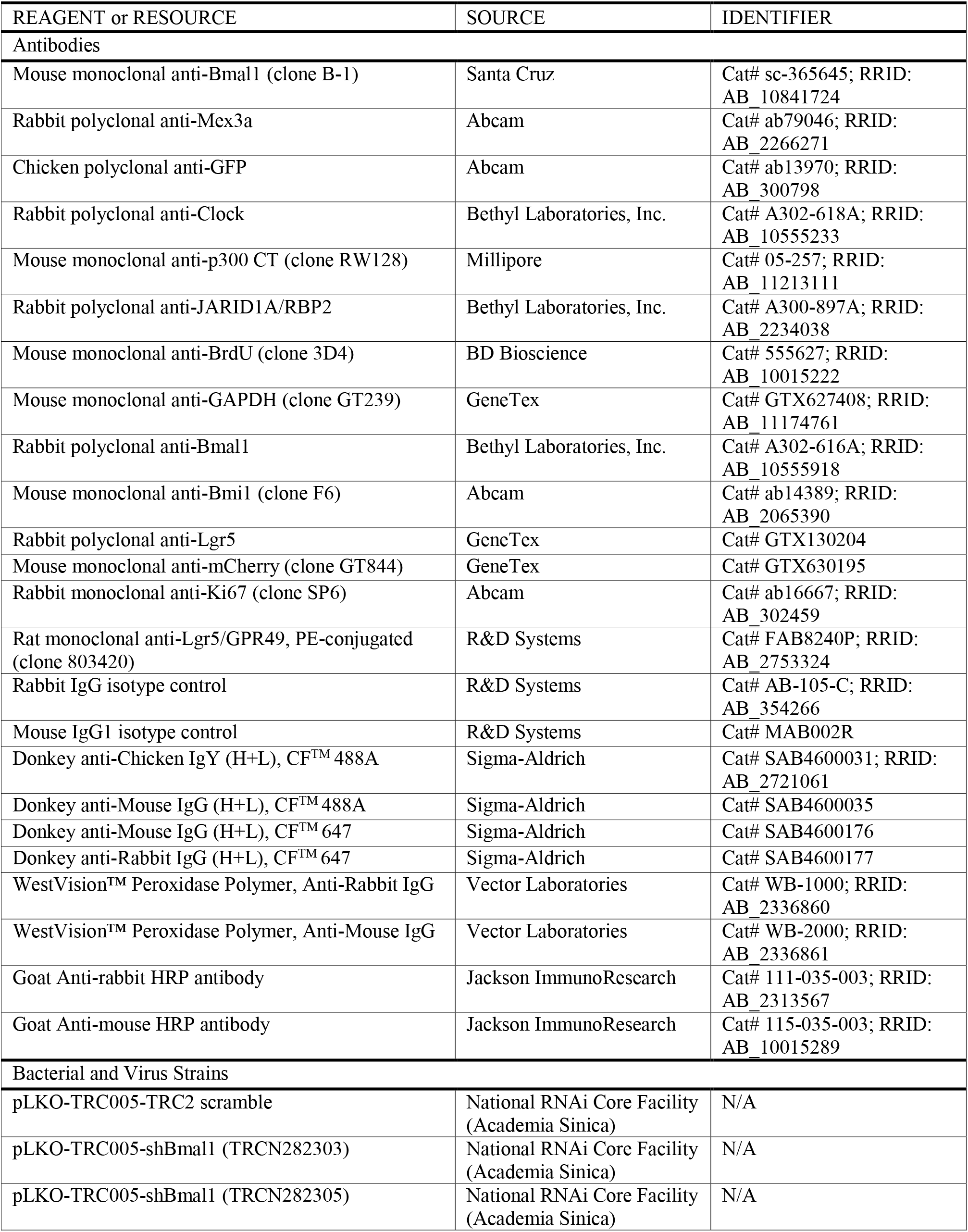
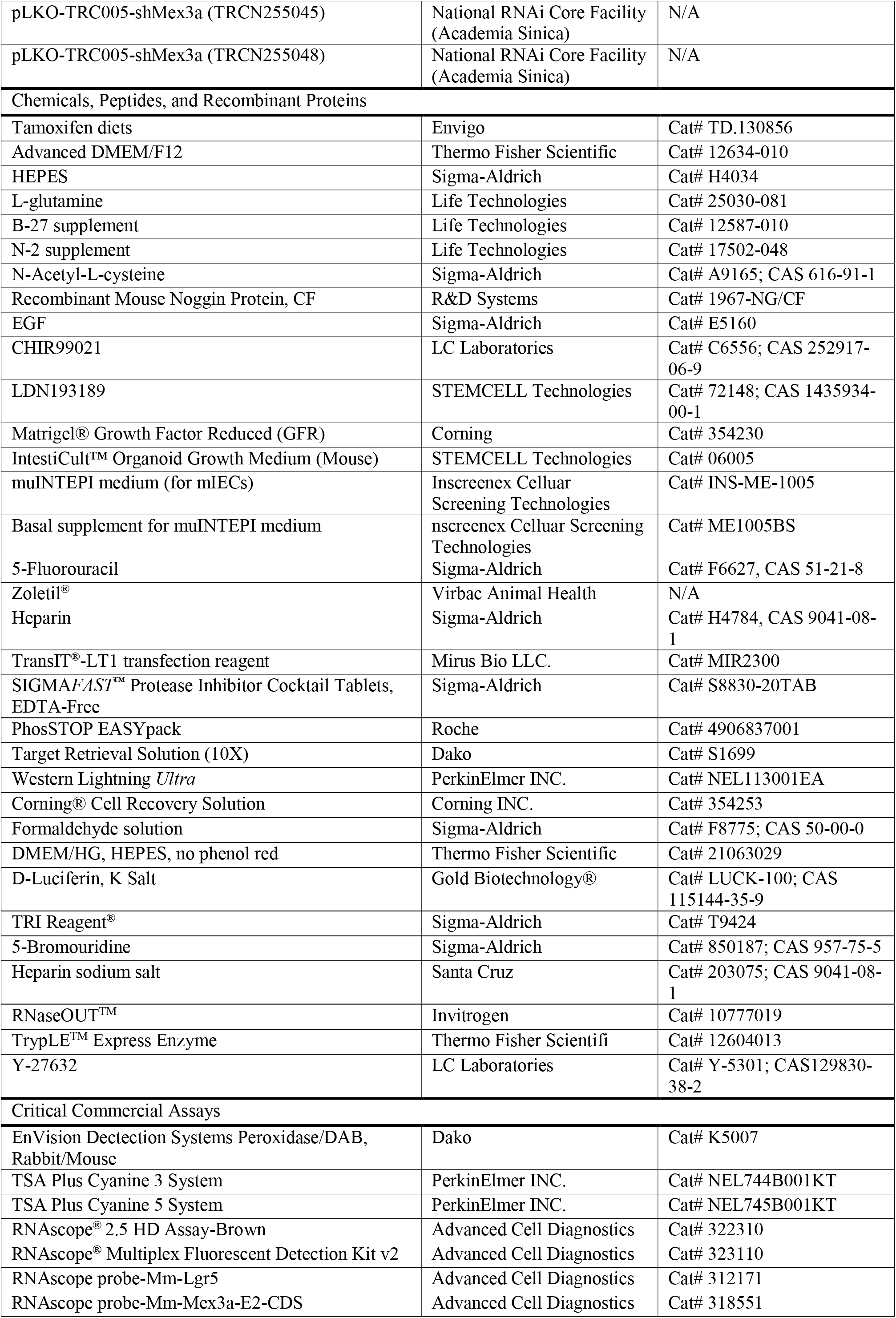
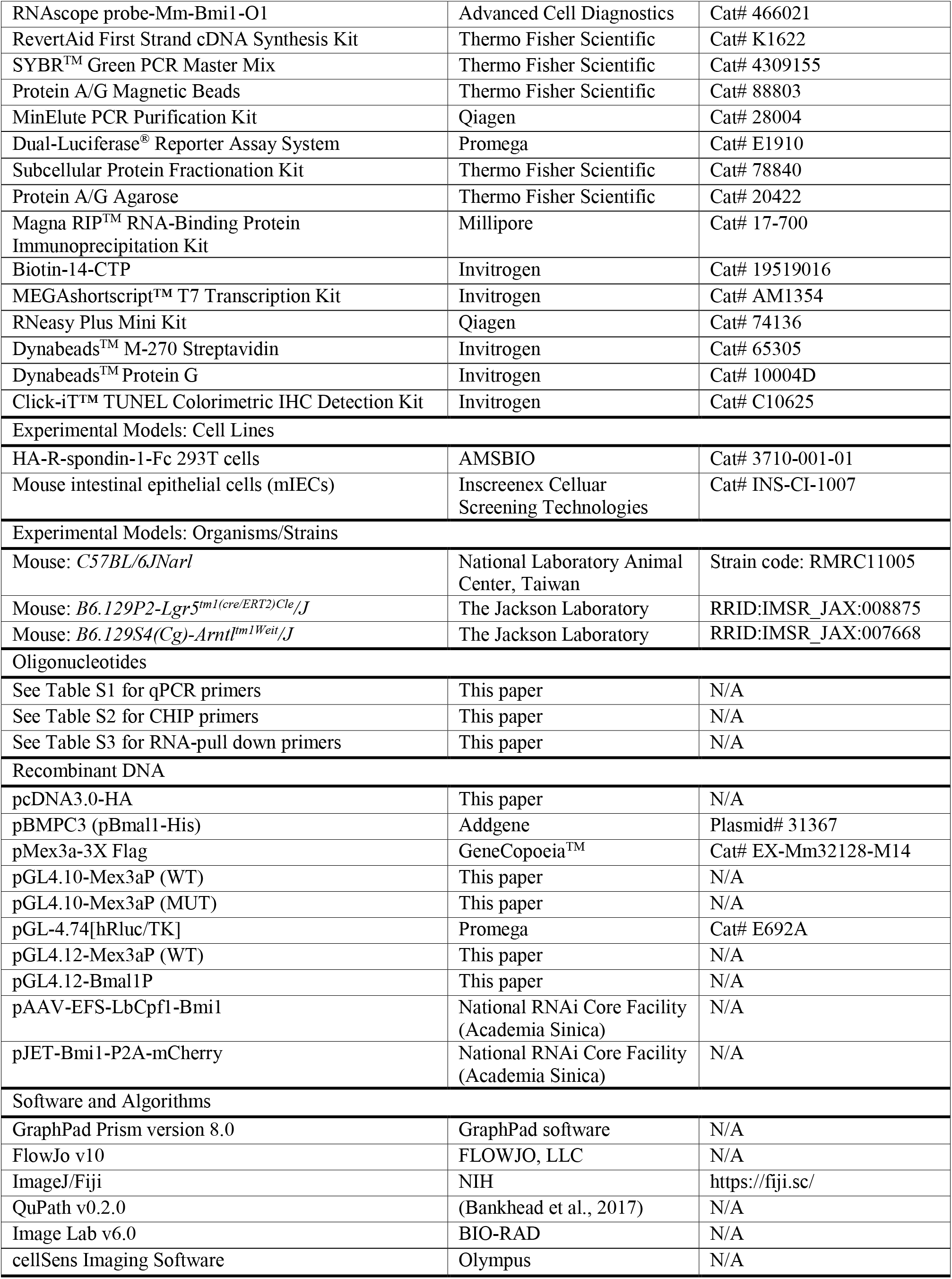

## CONTACT FOR REAGENT AND RESOURCE SHARING

Requests for further information, reagents, and resources should be directed to and will be fulfilled by the Lead Contact, Wendy W. Hwang-Verslues.

## EXPERIMENTAL MODEL AND SUBJECT DETAILS

### Mouse breeding and maintenance

All animal breeding and experimental procedures were approved by the Academia Sinica Institutional Animal Care and Utilization Committee (AS IACUC# 17-12-1165). All mice were maintained under standard 12:12 h light/dark (LD) condition, ZT0 (7:00 AM) corresponded to lights-on and ZT12 (7:00 PM) corresponded to lights-off. *C57BL/6JNarl* mice were purchased from the National Laboratory Animal Center (Taipei, Taiwan). To specifically knockout *Bmal1* expression in LGR5^+^ CBCs in the duodenum, *Lgr5-EGFP-ires-CreERT2* (referred as *Lgr5-Cre*; JAX stock no. 008875) mice (Barker et al., 2007) were crossed with *Bmal1* floxed mice with two *loxP* sites flanking exon8 of mouse *Bmal1* (*Bmal1^fl/fl^*; JAX stock no. 007668) (Storch et al., 2007). The progeny (*Lgr5-Cre^+^;Bmal1^fl/fl^, Lgr5-Cre^+^;Bmal1^wt/wt^* mice were used as control mice) were fed with tamoxifen diets (ENVIGO; TD130856) for 3 weeks to activate the CreERT2 recombinase (Figure 2A).

### Isolation of mouse intestinal crypt cells and organoid culture

Crypts were isolated from duodenum of *Lgr5-Cre^+^;Bmal1^wt/wt^* or *Lgr5-Cre^+^;Bmal1^fl/fl^* mice using the optimized protocol by Hans Clevers’ group (Sato et al., 2009). In brief, 5 cm duodenum was harvested and flushed by ice-cold phosphate buffered saline (PBS) to remove excrement. Tissues were cut into small pieces and the fragments were washed by ice-cold PBS for 10-20 times until the supernatant became clear. Fragments were then incubated in 20mL PBS/2mM EDTA with gentle agitation at 4°C for 20min. After removal of supernatant, the EDTA treated fragments were further disrupted mechanically in 10mL Advanced DMEM/F12 (Thermo Fisher Scientific, #12634-010). The supernatant was collected and filtered through 70μm cell strainers (BD Bio-sciences, Bedford, MA) as fraction 1. This process was repeated four times, and 2 fractions containing more intact crypts and less debris were combined. After centrifugation at 300 *g* for 3 min, crypt pellets were resuspended in a 1:1 mixture of organoid medium (Advanced DMEM/F12, 10mM HEPES, 2mM L-glutamine, 1X B27 supplement, 1X N2 supplement, 1mM N-acetyl-cysteine, 100ng/mL Noggin, 50ng/mL EGF, 250ng/mL R-spodin-1, 1μM CHIR99021, and 0.25μM LDN-193189) and growth factor reduced matrigel to reach 7 crypts/μL. Three drops of cell suspension (20μL each drop) was plated in each well of a non-coated 24-well plate and maintained in 500μL organoid medium/mouse IntestiCult Organoid Growth Medium at 37°C in a 5% CO2 humidified incubator. The medium was changed every other day.

### Intestinal epithelial cell culture

Small intestinal epithelial cell line (mIEC) from mouse E19.5 fetuses (Schwerk et al., 2013) were obtained from Insreenex Cellular Screening Technologies and maintained in muINTEPI medium at 37°C in a 5% CO2 humidified incubator.

### Jugular vein cannulation

Eight to twelve week old *Lgr5-Cre^+^, Lgr5-Cre^+^;Bmal1^wt/wt^* or *Lgr5-Cre^+^;Bmal1^fl/fl^* mice were used to perform the catheter implantation surgery as described previously (Kmiotek et al., 2012). Mice were anesthetized by intraperitoneal injection with the mixture of Zoletil (0.5mg/20g BW (body weight)) and Ropum (1.17mg/20g BW) and placed in the supine position under the dissecting microscope. After the neck area was sterilized with 70% alcohol, a 1cm long incision was cut from right clavicle to mandible of mice. The right jugular vein was isolated from the connective and adipose tissue under the skin of neck, and front 1cm of the 10cm 2 Fr PU tubing (Instech Laboratories, Inc BTPU-027) was implanted into the vein after a small incision was made on the jugular vein using micro scissors. Another side of PU tube was connected to a MOUSE VAB™ (Instech Laboratories, IncVABM1B/25), and the VAB was further placed under the skin of the back. After these mice recovered, PBS (control group) or 0.4mg/kg BW 5-FU was injected at ZT9 (4:00 PM) or ZT17 (00:00 AM) every other day. 100μL of 0.3mg/mL Heparin-solution was administrated to flush the catheters every day to prevent blood clots.

## METHOD DETAILS

### Immunoblot assay

mIEC were lysed in RIPA buffer (150mM NaCl, 1% NP-40, 0.5% sodium deoxycholate, 0.1% SDS, 50mM Tris pH8.0, SIGMAFAST™ Protease Inhibitors and PhosSTOP EASYpack phosphatase inhibitor cocktail). Protein concentration was determined by Bradford assay (Bio-Rad). 50-30 μg total proteins were separated by 7.5% or 10% tris-glycine polyacrylamide gel, with overnight incubation with primary antibodies (see the key resources table) and followed by a 1:10000 dilution of horseradish peroxidase (HRP)-conjugated secondary antibodies. HRP signals were detected using Western Lightning^®^ Ultra Chemiluminescence Substrate. The densitometry of blot bands was quantified using Image Lab 6.0 software.

### Immunohistochemical (IHC) and Immunofluorescence (IF) staining

Mice were entrained under LD condition for 3 weeks, then the duodenum tissues were harvested every 4 h starting from ZT1 (8:00 am) until ZT21 (4:00 am). Tissues were fixed in 10% formalin overnight at 4°C and embedded in paraffin. Sections were cut into 4 μm slices, dewaxed with xylene and rehydrated with descending ethanol series to water. Antigen retrieval was performed using the citrate-based antigen unmasking solution for 20 min under high pressure condition. Slices were stained with Rabbit anti-BMAL1 primary antibodies overnight at 4°C, followed by incubation with DAKO REAL™ EnVision™ HRP Rabbit/Mouse at room-temperature for 30 min. The 3.3’-diamiobenzidine (DAB) substrate was used to detect the peroxidase activity, and the slices were counterstained with hematoxylin. Slides were photographed under 40X magnification by the Aperio scanner machine (Leica Biosystems, Singapore). The percentage of positive cells was scored by the QuPath software.

For IF staining, tissues were hybridized with mouse anti-BMAL1 (1:50), rabbit anti-MEX3A (1:50) or chicken anti-GFP (1:500) primary antibodies overnight at 4°C after de-paraffined, rehydration and antigen retrieval. After washed by PBST three times, the slices were incubated with anti-mouse (1:1000), anti-rabbit polymer-HRP (1:1000) or anti-chicken CF488A (1:200) secondary antibodies at room-temperature for 1 h. To enhance the fluorescence signal, the anti-BMAL1 and anti-MEX3A hybridized slides were further labeled with TSA™ plus cyanine 3 (1:1500) or TSA™ plus cyanine 5 (1:1500), respectively. Images were obtained by a laser scanning confocal microscope (LSM770, Carl Zeiss MicroImaging). The fluorescence intensity and the length of crypt and villus were quantified using the QuPath and the CellSens software, respectively.

For organoid staining, the Corning^®^ Cell Recovery Solution was used to depolymerize the matrigel at 4°C for 30 min. The organoids were fixed in 4% paraformaldehyde in PME buffer (50mM PIPES, 2.5mM MgCl2, and 5mM EDTA) at room-temperature for 20 min. After permeabilization with 0.5% Triton X-100 for 20 min, the organoids were blocked with 5% BSA for 1 h. Antibody hybridization was performed as described above.

### RNAscope in situ hybridization

*Lgr5*, *Mex3a*, or *Bmi1* mRNA expression were detected by the RNAScope^®^ 2.5 HD Detection Reagent-Brown or RNAScope^®^ Multiplex Fluorescence Detection Reagent V2 kit with Mm-Lgr5, Mm-Mex3a-E2-CDS, and Mm-Bmi1-O1 probes according to the manufacturer’s instructions. Quantification of mRNA positive spots were performed using the trainable Weka segmentation classifier in Fiji software according to the recommendation of TECHNICAL NOTE from Advanced Cell Diagnostics.

### Quantitative real-time PCR

RNA was extracted using TRI Reagent according to the manufacturer’s instructions. For mRNA expression detection, 3 μg total RNA from each sample was reverse-transcribed using RevertAid First Strand cDNA Synthesis Kit. 10 ng 1st-strand cDNA was used for real-time PCR with appropriate primer sets and SYBR® Green PCR Master Mix using ABI-7900 thermocycler. Beta-glucuonidase (GusB) was used as an internal control. All primers are listed in Table S1.

### Chromatin immunoprecipitation (ChIP) assay

Cells were crosslinked with 1% formaldehyde for 10 min at 37°C and the crosslinking was stopped by 0.125 M glycine. After PBS wash, the crosslinked cells were lysed in 1 mL cell lysis buffer (10mM Tris-HCL pH8.0, 1mM EDTA, 0.5% NP-40, 1mM PMSF, SIGMAFAST™ protease inhibitor cocktail, and Roche PhoStop) on ice for 20 min. Nuclei pellet was then resuspended and incubated in 1 mL of nuclear lysis buffer (50mM Tris-HCL pH8.0, 100mM EDTA, 1% SDS, 1mM PMSF, SIGMAFAST™ protease inhibitor cocktail, and Roche PhoStop) on ice for 30 min and 3 freeze-thaw cycles to release chromatin. After centrifugation, the crosslinked chromatin was resuspended in 80 μL nuclear lysis buffers and sonicated using a Diagenode Bioruptor^®^pico (15 cycles of 30 sec on and 30 sec off) to 200~500 bases. A 10 μL aliquot of the fragmented chromatin was used as an input control. For immunoprecipitation assay, pre-washed protein A/G magnetic beads were incubated with 4 μg mouse anti-Bmal1 antibody at 4°C overnight. 25 μg sonicated chromatin was diluted to a final volume of 1mL with dilution buffer (20 mM Tris-HCL pH 8.0, 150 mM NaCl, 1 mM EDTA, 0.01% SDS, 1% triton X-100), and pre-cleaned with pre-washed protein A/G magnetic beads and 10 μg mouse-IgG isotype control at 4°C overnight. The antibody-coupled beads were then incubated with the sonicated chromatin for 4 h at 4°C. After washed several times, the chromatin-protein complex was reversed crosslinked by addition of 15 μL solution containing 2.5 M NaCl, 2 μL 50mg/mL RNaseA and 6 μL 10 mg/mL proteinase K. DNA was extracted by MinELute^®^ PCR purification kit. Semi-quantitative PCR was performed to detect BMAL1 associated promoter regions using primers listed in Table S2.

### Luciferase reporter assay

Mus muscles strain *C57BL/6JNarl* chromosome 3 (88,530,849-88,533,098 bp) containing the proximal promoter of *Mex3a*, *Mex3a* exon1 and part of the coding sequence, was amplified by PCR from mIEC genomic DNA and cloned into pGL4.10 vector (referred as pGL4.10-Mex3aP (WT)). The putative Bmal1 binding site predicted by the JASPAR database was mutated from 5’-GCCTTTCCAC-3’ to 5’-TTTTTTTTAT-3’ (referred as pGL4.10-Mex3aP (MUT)). pGL4.74 Renilla luciferase plasmid was used as a transfection control. For HEK-293T cells, 6×10^5^ cells were seeded and co-transfected with 500ng of pGL4.10-Mex3aP (WT) or pGL4.10-Mex3aP (MUT), 6.25 ng of GL4.74, pcDNA3.0 or pcDNA3.0-Bmal1-His (range from 300 ng to 600 ng) using TransIT-LT1 transfection reagents. Cell extracts were collected 48 h after transfection and the luciferase activity was measured using Dual-Luciferase® Reporter (DLR™) Assay System.

### Co-immunoprecipitation (Co-IP) assay

Nuclear extracts were collected using Subcellular Protein Fractionation Kit according to the manufacturer's instruction. For IP, 300 μg nuclear lysate was incubated with 4 μg anti-BMAL1, 9 μg anti-CLOCK, 9 μg anti-P300, 6 μg anti-JARID1A, or control IgG antibody at 4℃ overnight. The antibody-protein complexes were precipitated using 20 μL pre-washed and pre-blocked (10% BSA) protein A/G agarose beads at 4℃ for 1 h. After washed with NP-40 TNE buffer (10mM Tris-HCl pH7.5, 150mM NaCl, 0.5mM EDTA, 0.1% NP-40, and protease inhibitor cocktail) 5 times, interacting proteins were eluted with SDS-PAGE and detected by western blotting assay.

### RNA immunoprecipitation (RIP) assay

Whole-cell lysate from 2 × 10^7^ mIEC was used for RIP assay using Magna RIP™ RNA-Binding Protein Immunoprecipitation Kit according to the manufacturer's instruction. 4 μg anti-MEX3A antibody was used for the immunoprecipitation at 4℃ overnight, and rabbit IgG was used as a pull-down control. MEX3A bound RNAs were then isolated following the procedures of proteinase K digestion and phenol-chloroform-isoamyl alcohol extraction. The precipitated RNAs were resuspended in 12 μL DEPC-ddH2O. Quantitative PCR was performed to determine the level of MEX3A-bound *lgr5* or *bmi1* mRNA.

### RNA pull down assay

Biotin-14-CTP labeled RNA was *in vitro* transcribed using MEGAshortscript™ T7 High Yield Transcription Kit according to the manufacturer's instruction. After purification with RNeasy Plus Mini Kit, 2 μg biotinylated RNAs was incubated with 20 μL pre-washed Dynabeads™ M-270 Streptavidin in 1 mL buffer (20 mM Tris-HCl (pH7.5), 100 mM KCl, 5 mM MgCl_2_, and 0.5% NP-40), and incubated at 4°C for 1 h with gentle agitation. 450 μg mIEC cytoplasmic protein lysate was added to the washed RNA-probe coupled dynabeads, and incubated in 1× TENT buffer (10 mM Tris-HCl (pH8.0), 1 mM EDTA (pH 8.0), 250 mM NaCl, and 0.5% Triton X-100) with SIGMAFAST™ proteinase inhibitor cocktails and RNaseOUT™ Recombinant Ribonuclease Inhibitor at 4°C for 8 h. After washed with 1× TENT buffer, the proteins interacted with mouse *lgr5* probes were eluted using 40 μL SDS buffer and detected by immunoblot assay. Biotinylated *Gapdh* RNA was used as a pull-down control.

### BRIC assay

mIEC was transfected with pcDNA3.0 or pMex3a-3X Flag and then incubated in the 5’-bromo-uridine (BrU) containing medium to a final concentration of 1mM in a humidified incubator with 5% CO2 for 8 h. After removal of BrU, RNA was harvested at indicated time points using the TRI Reagent^®^. 15 μL pre-washed Dynabeads™ Protein G was blocked with 2% BSA and 5 mg heparin at 4°C for 1 h, and then incubated with 4 μg anti-BrdU Ab at 4°C for 1 h. The BrdU Ab coupled dynabeads were further washed using BrU-IP buffer and 250 mM NaCl three times and resuspended in 15 μL BrU-IP buffer supplemented with RNaseOUT™ Recombinant Ribonuclease Inhibitor. To normalize the immunoprecipitation of BrU-labeled RNAs among different time points, the *in vitro* transcribed BrU-labeled luciferase RNA was used as a spike-in control for internal standard of quantitative PCR assay. 50 μg BrU-labeled RNAs from mIEC and 100 ng BrU-labeled luciferase RNA were mixed in 1 mL BrU-IP buffer with RNaseOUT, and the RNA mixture was denatured by heating at 80°C for 3 min. Subsequently, 15 μL BrdU Ab conjugated dynabeads were added into RNA mixture and incubated at room temperature for 1 h. After washed four times with BrU-IP buffer (20 mM Tris-HCL pH7.5, 250 mM NaCl, and 20 μL RNaseOUT), the BrU-labeled RNAs were isolated using RNeasy Plus Mini Kit. The half-life of *lgr5* or *bmi1* mRNA was determined by RT-qPCR assay.

### Generation of CRISPR-mBmi1-P2A-mCherry plasmid, electroporation and generation of knock-in cells

To track cell lineage of BMI1-expressing cells, an all-in-one AAV plasmid (pAAV-EFS-LbCpf1-Bmi1) was constructed. The crRNA of LbCpf1 and the *Bmi1* sgRNA (5’-cttaacagtcctaaccagatgaag-3’) to target 3’-end of coding sequence at *Bmi1* exon10 were transcribed under the control of U6 promoter, and the elongation factor-1α short (EFS) promoter drives the LbCpf1 expression. 1.5 μg pAAV-EFS-LbCpf1-Bmi1 and 1.5 μg *Bmi1* exon10-encoding and P2A-mCherry-encoding PCR amplicons were simultaneously transfected into 2 × 10^5^ mIEC using TransIT-LT1 transfection reagents. After enrichment of the BMI1^+^ cell populations by sorting out the mCherry^high^ mIEC, the double immunofluorescence staining with BMI1 and mCherry was performed to validate the BMI1 expression in P2A-mCherry knock-in mIEC.

### FACS sorting and analysis

Duodenum organoids from *Lgr5-Cre^+^;Bmal1^wt/wt^* or *Lgr5-Cre^+^;Bmal1^fl/fl^* mice were collected and dissociated by pipetting. The dissociated cell aggregates were further incubated in TrypLE with 100 μM Y-27632 at 37°C for 10 min to separate to single cells. Cells were resuspended in organoid medium containing 100 μM Y-27632 and filtered through a 70 μm strainer (BD Bioscience). A broad FSC-A/FSC-W and SSC-A/SSC-W was gated as single cell population, and the LGR5-EGFP^+^ cells were sorted.

Bmi1-P2A-mCherry knock-in mIEC and parental mIEC were trypsinized and then resuspended in 100 μM Y-27632 containing medium. 5×10^6^ mIEC were further labeled with 1 μg anti-mouse Lgr5/GPR49 PE-conjugated antibodies in 100 μL medium at room temperature for 1 h. Both Bmi1-P2A-mCherry knock-in or LGR5 stained mIECs were filtered through a 70 μm strainer. PE^+^ or PE-Texas Red^High^ were sorted as LGR5^+^ or BMI1^high^ cells using the BD FACS Aria II cell sorter, respectively.

### Quantification and statistical analysis

Quantitative results were represented as Mean ± SD. Statistical analysis was conducted using the Graph Pad Prism 8.0 (GraphPad Software). Non-parametric Mann-Whitney test was used to compare control and experimental genotypes or treatment groups. Statistical significance among more than three groups was analyzed using one-way ANOVA with Dunn’s multiple comparison test. For the time course analysis of mRNA expression levels in organoids with tamoxifen treatment and BRIC assay, two-way ANOVA with Sidak multiple comparison test was performed to detect significance among different groups. Asterisk (*, **, ***) indicates statistical significance with *p*-value <0.05, 0.01, 0.001, respectively.

## SUPPLEMENTAL INFORMATION

Supplemental Information can be found online at ~~

## ACKNOWLEDGEMENTS

This work was supported by Academia Sinica [AS-SUMMIT-109] and Ministry of Science and Technology [AS-KPQ-109-BioMed] [MOST 109-0210-01-18-02]. We thank Drs. Kuo-I Lin, Muh-Hwa Yang and Ruey-Hwa Chen for discussion. We thank the National RNAi Core Facility at Biomedical Translation Research Center, Academia Sinica, Taiwan for generating the mBmi1-P2A-mCherry CRISPR-related plasmids and supporting for lentivirus packaging; the flow cytometry and histology core facilities at Genomics Research Center, Academia Sinica for technical support; and the DNA sequencing core facility at IBMS Academia Sinica for providing DNA analysis. We also thank Academia Sinica Advanced Optics Microscope Core Facility for microscope imaging technical support. The core facility is funded by Academia Sinica Core Facility and Innovative Instrument Project (AS-CFII-108-116).

## AUTHOR CONTRIBUTIONS

W.W.H.-V. and F.-P.C. conceived and designed the studies. F.-P.C. performed experiments. C.-K.W. performed co-IP experiments of BMAL1 and other cofactors. T.-J.C. repeated the BMAL1 protein oscillation experiments using B6 WT mice. Y.-C.C established the Bmi1-P2A-mCherry lineage tracing mIEC cell line. P.-H.H. performed LC-MS/MS analysis in the beginning stage of this study. S.-S and Y.-W.W. maintained the mice and related experiments used in this study. F.-P.C. analyzed the data. W.W.H.-V. and F.-P.C. wrote the manuscript, and W.W.H.-V. supervised this study.

## DECLARATION OF INTERESTS

The authors declare no competing interests.

## Notes

**Conflict of Interest:** No direct competing interest.

### Competing Interest Statement

The authors have declared no competing interest.

