## Supplementary Figure and Tables for "BMAL1 and MEX3A co-regulate intestinal stem cell succession"

**This file includes:**

**Figures S1.** Bmal1 knockout in $LGR5^+$ CBCs alters expression of lineage specific genes in duodenum. (Related to Figure 2)

**Figure S2.** BMAL1 and MEX3A contribute to Bmi1 suppression. (Related to Figure 4)

**Figure S3.** Establishment of Bmi1-P2A-mCherry mIECs using CRISPR-Cpf1. (Related to Figure 6)

**Figure S4.** Proposed model. (Related to Discussion)

**Table S1.** Primer sequence for qPCR, related to STAR methods.

**Table S2.** Primer sequence for ChIP analysis, related to STAR methods.

**Figure S1** (related to Figure. 2)

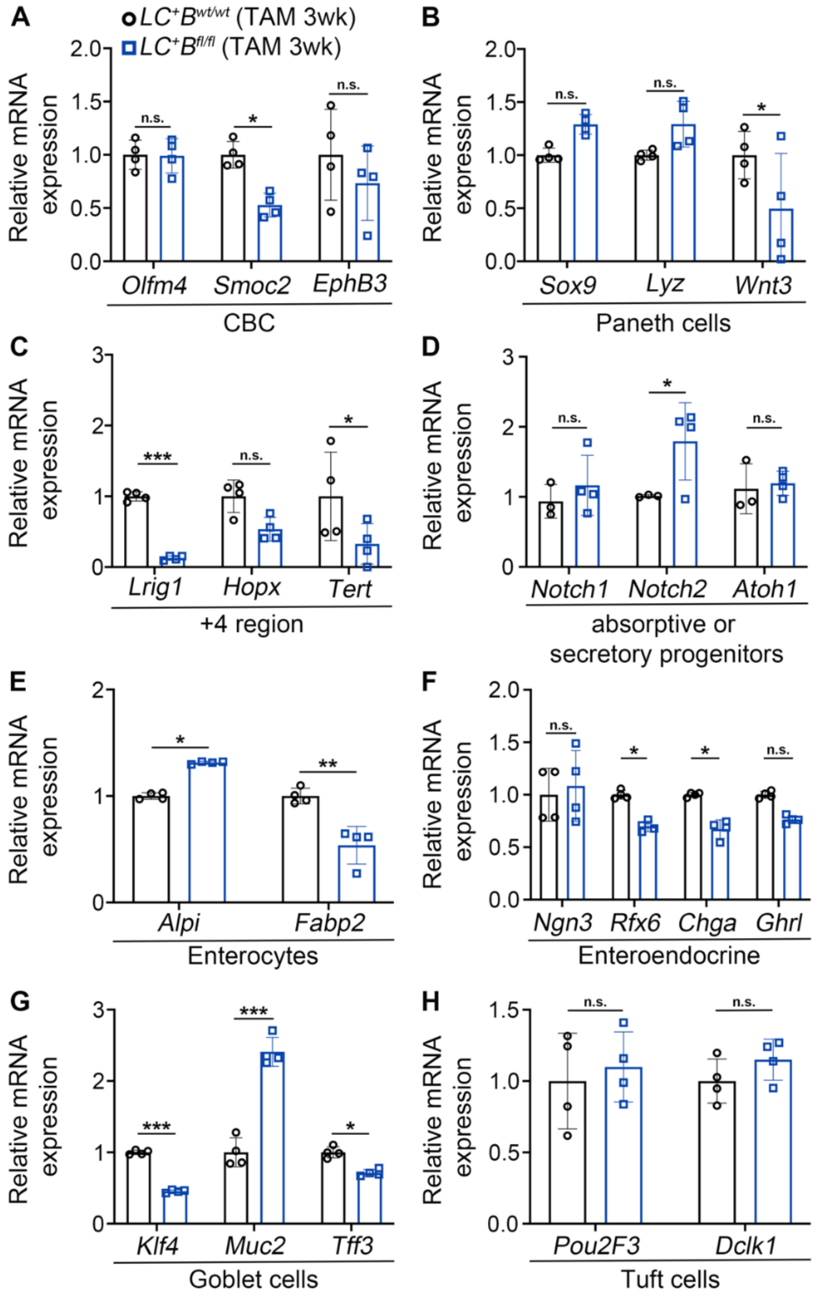


**Figure S1. Bmal1 knockout in LGR5$^{+}$ CBCs alters expression of lineage specific genes in duodenum.** (Related to Figure 2)

RT-qPCR of lineage specific genes in CBCs (A), Paneth cells (B), progenitors in +4 region (C), absorptive or secretory progenitors (D), enterocytes (E), Enteroendocrine (F), Goblet (G) and Tuft cells (H) from duodenum epithelium of TAM fed *Lgr5-Cre*$^{+}$*;Bmal1*$^{wt/wt}$ (*LC*$^{+}$*B*$^{wt/wt}$) control and *Lgr5-Cre*$^{+}$*;Bmal1*$^{fl/fl}$ (*LC*$^{+}$*B*$^{fl/fl}$) mice. Duodenum was harvested at ZT5. Data are presented as means ± SD, significant difference are based on Student's t test. * $P < 0.05$; ** $P < 0.01$; *** $P < 0.001$; n.s., no statistically significant difference.

**Figure S2** (related to Figure 4)

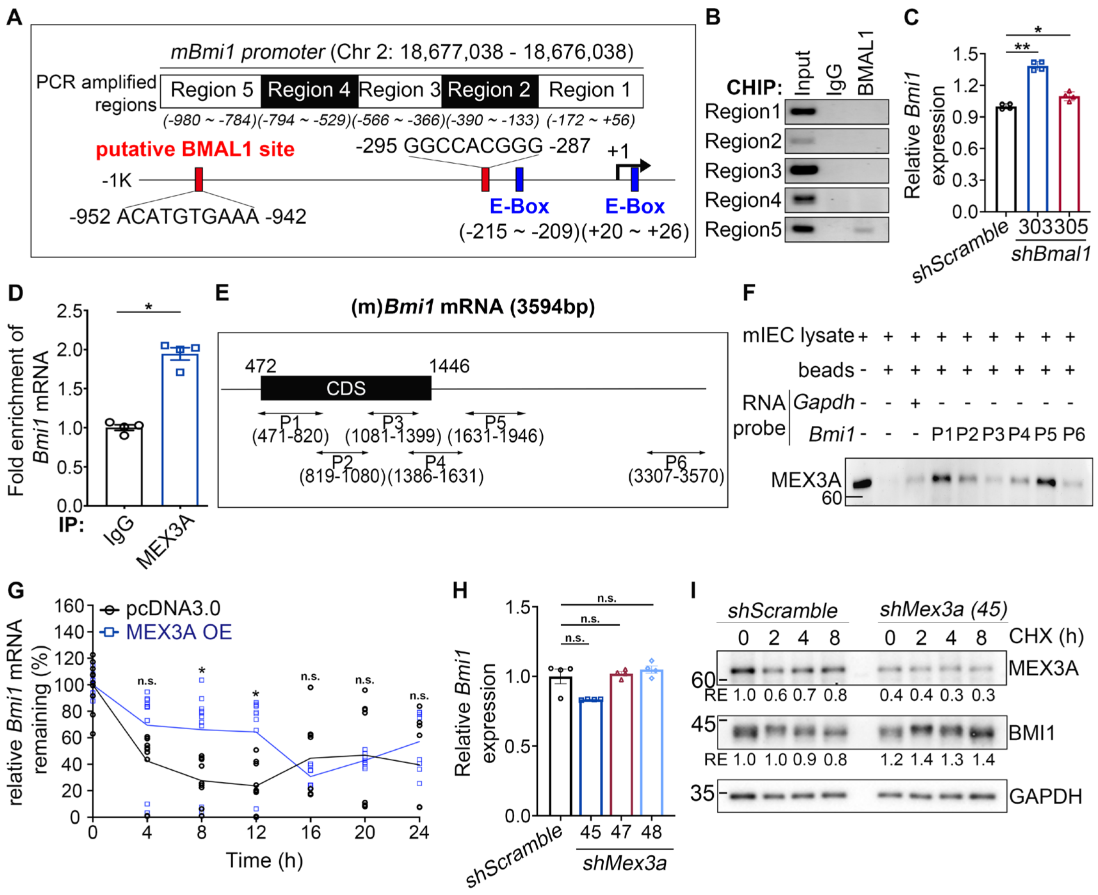


**Figure S2. BMAL1 and MEX3A contribute to *Bmi1* suppression.** (Related to Figure 4)

(A) Diagram shows two putative BMAL1 binding sites (red box) and two E-boxes (blue box) on the mouse *Bmi1* promoter.

(B) ChIP analysis of BMAL1 occupancy on the *Bmi1* promoter in mIECs. IgG was used as a negative control. Gels shown are from one representative experiment of two independent experiments.

(C) qRT-PCR analysis of *Bmi1* in mIECs transduced with *shScramble* control or *shBmal1* lentiviruses (clone #303 or #305). Three independent experiments were performed. Data show mean ± SD, significant differences are based on one-way ANOVA; * $P < 0.05$, ** $P < 0.01$.

(D) RNA-IP of MEX3A-*Bmi1* complex using mIEC lysates and anti-MEX3A antibody. IgG was used as a negative control. Three independent experiments were performed. Data are presented as mean ± SD, significant differences are based on Student's t-test; * $P < 0.05$.

(E) Diagram of mouse *Bmi1* mRNA and six biotinylated RNA probes (P1 to P6) spanning nearly whole Bmi1 mRNA sequence used for RNA pull-down assay.
(F) Immunoblot of MEX3A from *Bmi1* RNA pull-down using mIEC lysates with biotinylated RNA probes. Beads only and the *Gapdh* RNA probe were used as controls. This experiment was performed two times with similar results.
(G) BRIC of *Bmi1* mRNA in mIECs transfected with pcDNA3.0 control vector or pMex3a-3X flag *Mex3a* level at 0 h was set as 100%. Three independent experiments were performed. Data are represented as mean ± SD based on two-way ANOVA; * $p < 0.01$, n.s., no statistically significant difference.
(H) qRT-PCR of *Bmi1* levels in mIECs transduced with *shScramble* control or *shMex3a* lentiviruses (clone #45, #47 or #48). Three independent experiments were performed. Data show mean ± SD, significant differences are based on one-way ANOVA; n.s., no statistically significant difference.
(I) Immunoblot of BMI and MEX3A in cycloheximide (CHX) treated mIECs transduced with *shScramble* control or *shMex3a* lentiviruses. Blots shown are from one representative experiment of two independent experiments. RE: relative expression.

**Figure S3** (related to Figure 6)

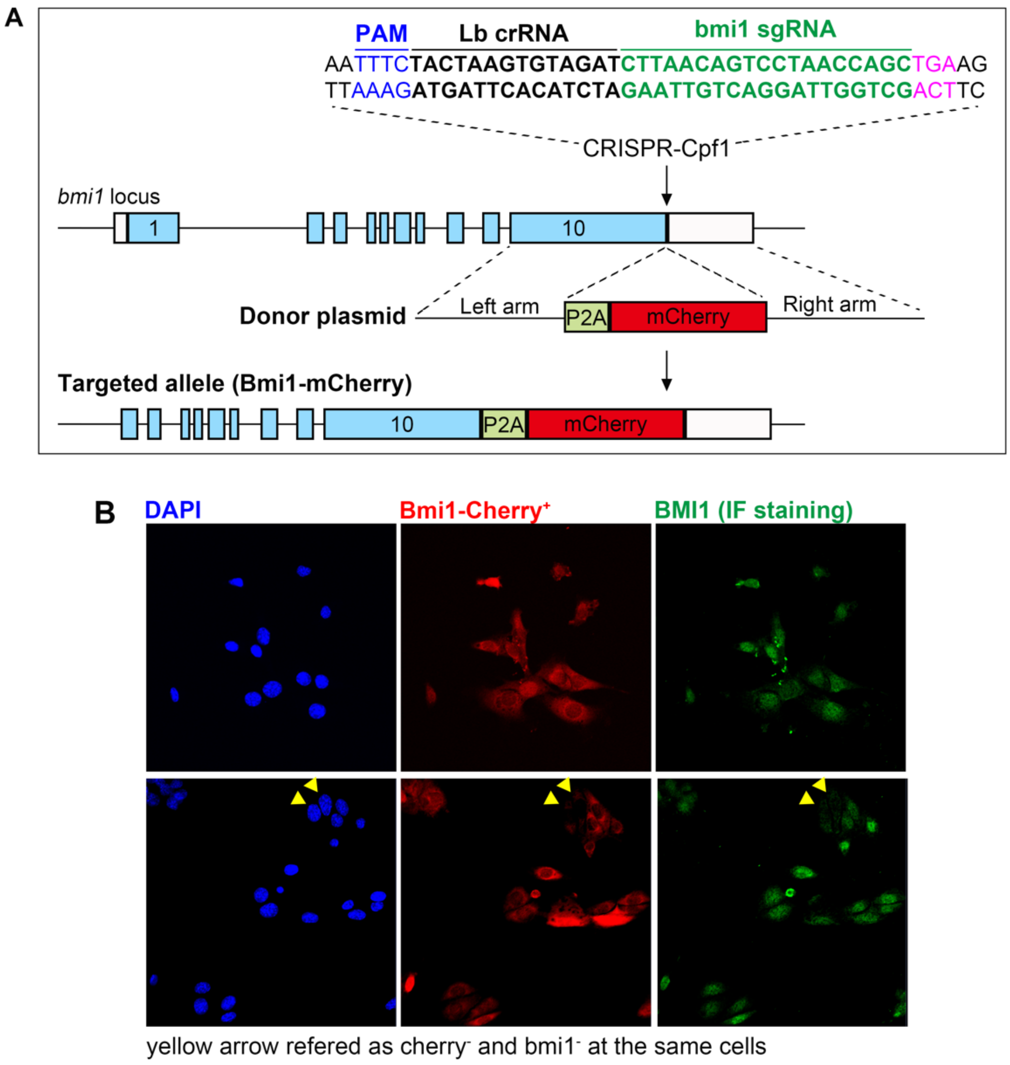


**Figure S3. Establishment of Bmi1-P2A-mCherry mIECs using CRISPR-Cpf1.** (Related to Figure 6)

(A) Diagram shows the insertion strategy of P2A-tdTOMATO behind exon10 of mouse *Bmi1* genes. PAM, crRNA and sgRNA in Cpf1 expression cassette are labeled with blue, black, and green lines, respectively.

(B) Immunofluorescence staining of BMI1 in sorted Bmi1-P2A-mCherry lineage traced mIECs. Red indicates the mCherry$^{+}$ cells. Yellow arrows indicate mCherry$^{-}$ mIECs without BMI1 expression.

**Figure S4** (related to Discussion)

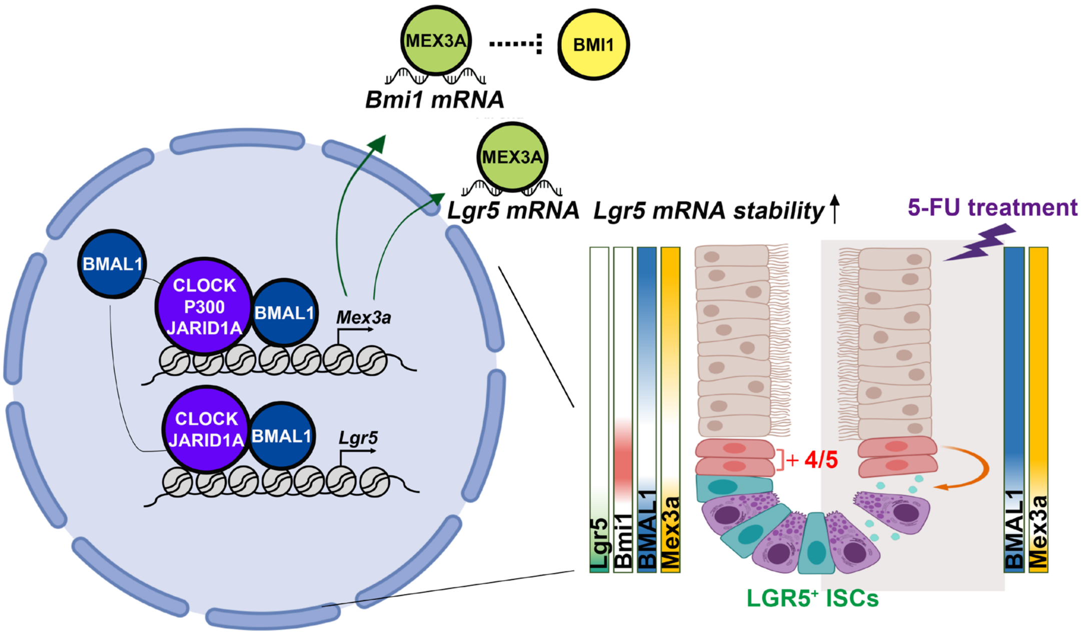


**Figure S4. Proposed model.** (Related to Discussion)
In intestinal crypts, BMAL1 and MEX3A ultradian oscillation is crucial to maintain homeostasis between the fast-proliferating LGR5$^{+}$ CBCs and damage-resistant BMI1$^{+}$ +4 cells. Ablation of *Bmal1* reduced both LGR5$^{+}$ CBCs and MEX3A expression but increased BMI1 levels. Induction of BMAL1 and MEX3A was sufficient to promote reprogramming of BMI1$^{+}$ cells to express LGR5. BMAL1, together with CLOCK and JARID1A, upregulated expression of both *Lgr5* and *Mex3a* and while Mex3a further enhanced Lgr5 expression by directly binding and stabilizing *Lgr5* mRNA. Based on this mechanistic insight, 5-FU delivery at ZT17 when BMAL1, Mex3a and Bmi1 level were high protects crypt cells from apoptosis and allows sustained ISC cell proliferation.

**Table S1. Primer sequence for qPCR, related to STAR methods.**

| Gene | Forward Primer | Reverse Primer |
|---|---|---|
| *Bmal1* | TGCCCTCTGGAGAAGGTGG | GGGGAGGCGTACTTGTGATG |
| *Mex3a* | GCAGGCAAGGCTGCAAGATT | ACTTGTTGCGTGAGGCTCTT |
| *Lgr5* | CCTTCCCCAGGTCCCTTCAA | GAACACGGTCAAAGCCACCA |
| *Bmi1* | CCGCTCTTTCCGGGATCTTT | GGACAATACTTGCTGGTCTCCA |
| *Olfm4* | CCTGCTCCTGGAAGCTGTAG | CAAAGGTGCCACCCAGTACA |
| *Smoc2* | GCAGGGAAAGCAGATGATGC | TGGTCTTGTTCTGCCGACTC |
| *EphB3* | ACTGCAGAAGATCTGCTAAGGA | CGTTGGAGCTGAGTGTCAGA |
| *Sox9* | CTGAAGGGCTACGACTGGAC | TTCAGCAGCCTCCAGAGCTT |
| *Lyz* | AAGGTCTACAATCGTTGTGAGTTG | GCTGAGCTAAACACACCCAGTC |
| *Wnt3* | TCCAACTATTGGGGGCGTCG | GGGCCAGGGACCACCAAA |
| *Lrig1* | TTCTGCTCTGGCTGCTCTTG | CAGGTTTAGGCTGCGCGTC |
| *Hopx* | GCGACTTTCAGTGGTTCCTGC | TCCGTGCGCGTGTGGAAG |
| *Tert* | CTGGCAGGGAAGGACCGTG | CCGTAGCCGCACTCTCTCAA |
| *Notch1* | TCAGTGCCCCAAAGGCTTCA | CATAGTGGCAGGGGTCAGGG |
| *Notch2* | CCCTTGCTTGAACGATGGGC | TTGCACCTCTGGCCTGTGAA |
| *Atoh1* | CGGGGCTTATCCCCTTCGTT | TCGGTGCTATCCAGGAGGGA |
| *Alpi* | CTCTGTGGGGTCAAGGCCAA | CTCCCACAGACTTCCCTGCT |
| *Fabp2* | CGGCACGTGGAAAGTAGACC | GGTCCAGGCCCCAGTGAG |
| *Ngn3* | TCGGGAGAACTAGGATGGCG | TCGGCAGTCACCCACTTCTG |
| *Rfx6* | GCGGTGGGAAGGATGGCTAA | TGCATTTCTGATTTAACTCCCACC T |
| *Chga* | CAAGCACAGAGACGCAGCAG | ATTGGTGGCTGTGTCCTCCC |
| *Ghrl* | AAGGAGAAGCCGGTGAGCAG | TTCTCTGCTGGGCTTTCTGGT |
| *Klf4* | CTGCGAACTCACACAGGCGA | AGGCCCTGTCACACTTCTGG |
| *Muc2* | GGGACAAGTGTGGCTGCTAT | TGAAGTGTTGCCCCACTGTT |
| *Tff3* | TACGTTGGCCTGTCTCCAAG | TGCAGAGGTTTGAAGCACCA |
| *Pou2f3* | GGAGCCCATGCACACAGAGA | GGAGAGAAGCCATGTCCCCAG |
| *Dclk1* | GCATCCCTGGGTTAATGATGATGG | GAAACTCCTGCTGCAGTGCT |
| *GusB* | CCGACCTCTCGAACAACCG | GCTTCCCGTTCATACCACACC |

**Table S2. Primer sequence for ChIP analysis, related to STAR methods.**

| Promoter/Site | Forward Primer | Reverse Primer |
|---|---|---|
| *Mex3a*/Region1 (-830~-403) | GTCTGCTCCACCAGCTCTCT | CTTCCAGCGCTCAAACTCTT |
| *Mex3a*/Region2 (-1111~-831) | TGGCTTCCCGGATCAGCCCC | AAAGCCCGCGGCCGCTACAG |
| *Mex3a*/FSC (far site control) | CCCAGGAGTCCCCCACTTAC | TGGCACTTCTCTAGCCCTGC |
| *Lgr5*/Region1 (-2404~-2001) | GTCTGGGAGGCAGCTCCAAA | TGCTTGATCGCAGACCCCAA |
| *Lgr5*/Region2 (-2020~-1569) | TTGGGGTCTGCGATCAAGCA | CACAATACACCCACTGCGCC |
| *Lgr5*/Region3 (-1568~-1063) | AATGAGAGAGGAGGTAAGGGA AAGA | CAAGTCCACCTAGATTTGATTC TTAG |
| *Lgr5*/Region4 (-1062~-694) | AATGCCCCAAAGGGCTACCGG A | AACCAAGAGCCACCCCACTC |
| *Lgr5*/Region5 (-713~-270) | GAGTGGGGTGGCTCTTGGTT | GGTACCCGCGACTGAGATGT |
| *Lgr5*/Region6 (-269~+25) | CAGGAACTGTAGAGGATGGGT GCAG | TGCTGCAGGCGGTCTGAGCTGT GTG |
| *Lgr5*/FSC (far site control) | ACCCCCGTGACACGTTTTCT | GGGTGGGGGTGGGAAAAAGA |
